# Predicting Minimal Residual Disease in Acute Myeloid Leukemia through Stochastic Modeling of Clonality

**DOI:** 10.1101/790261

**Authors:** Khanh Dinh, Roman Jaksik, Seth J. Corey, Marek Kimmel

**Affiliations:** Department of Statistics, Rice University, Houston, TX, USA; Systems Engineering Group, Silesian University of Technology, Gliwice, Poland; Departments of Pediatric Hematology/Oncology and Stem Cell Transplantation and Cancer Biology, Cleveland Clinic, Cleveland, OH, USA; Departments of Statistics and Bioengineering, Rice University, Houston, TX, USA Systems Engineering Group, Silesian University of Technology, Gliwice, Poland

## Abstract

Event-free and overall survival remains poor for acute myeloid leukemia (AML). Chemo-resistant clones contributing to relapse of the disease arise from minimal residual disease (MRD) rather than resulting from newly acquired mutations during or after chemotherapy. MRD is the presence of measurable leukemic cells using non-morphologic assays. It is considered a strong predictor of relapse. The dynamics of clones comprising MRD is poorly understood and is considered influenced by a form of Darwinian selection. We propose a stochastic model based on a multitype (multi-clone) age-dependent Markov branching process to study how random events in MRD contribute to the heterogeneity in response to treatment in a cohort of six patients from The Cancer Genome Atlas database with whole genome sequencing data at two time points. Our model offers a more accurate understanding of how relapse arises and which properties allow a leukemic clone to thrive in the Darwinian competition among leukemic and normal hematopoietic clones. The model suggests a quantitative relationship between MRD and time to relapse and therefore may aid clinicians in determining when and how to implement treatment changes to postpone or prevent the time to relapse.

**Author summary:** Relapse affects about 50% of AML patients who achieved remission after treatment, and the prognosis of relapsed AML is poor. Current evidence has shown that in many patients, mutations giving rise to relapse are already present at diagnosis and remain in small numbers in remission, defined as the minimal residual disease (MRD). We propose a mathematical model to analyze how MRD develops into relapse, and how random events in MRD may affect the patient’s fate. This work may aid clinicians in predicting the range of outcomes of chemotherapy, given mutational data at diagnosis. This can help in choosing treatment strategies that reduce the risk of relapse.

## Introduction

Acute myeloid leukemia (AML) is the most common myeloid malignancy with over 21,000 cases diagnosed annually in the United States. Event-free and overall survival remain poor, chiefly because of the emergence of chemoresistant clones. Despite stem cell transplantation and new drug approvals, chemoresistance and relapse remain major obstacles to survival for those with AML.

Different models have been proposed for the emergence of AML and its treatment outcomes as well as for other hematologic malignancies (1–10). Deterministic models, of which an example is the model of (2) do not account for the unpredictability of clonal heterogeneity in the genomic landscapes or therapeutic responses of an individual’s cancer. The main objective of the present work is to characterize the range of treatment outcomes by taking into account that when leukemic cell population is reduced to very low numbers, survival of any particular clone is subject to chance fluctuations. That is, even with the same parameters of disease state such as blast percentages in bone marrow and peripheral blood, leukemic clonal percentages at diagnosis, and treatment responses such as chemotherapy-induced cell death rate (see **Table 1** for a complete listing), a small residual population of malignant cells may or may not regrow. Another objective is to verify if the extent of the minimal residual disease (MRD) is a predictor of the time to relapse.

**Table 1:**
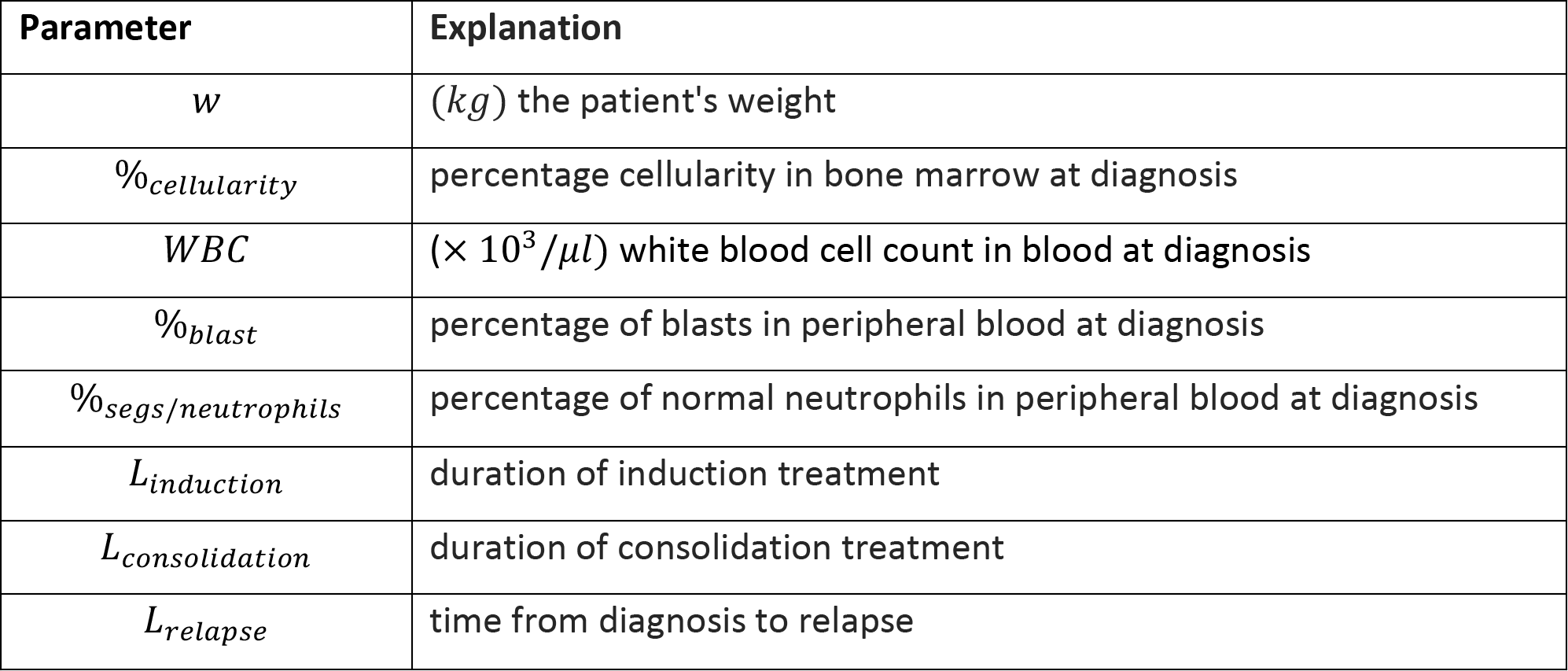
Data available in the TCGA used as inputs for modeling.

We developed a stochastic model of clonal evolution based on a multitype (multi-clone) age-dependent Markov branching process model of cell proliferation (11). In brief, we consider the critical time interval between diagnosis and initial relapse of AML that includes cytotoxic chemotherapy, chemotherapy-induced myelosuppression and decrease in leukemic cells, non-leukemic marrow recovery, and growth of the leukemic clones due to refractory or relapsed disease. **Figure 1** depicts a simplified sequence of events we address (including, for completeness, leukemogenesis that we do not model). This scenario may be different if chemotherapy leads to eradication of the malignant cells, i.e., absence of minimal residual disease (MRD). However, the most informative data at are those from patients suffering relapse with the malignant myeloid bone marrow cells being sequenced at two time points: diagnosis and at relapse.

**Fig 1.**
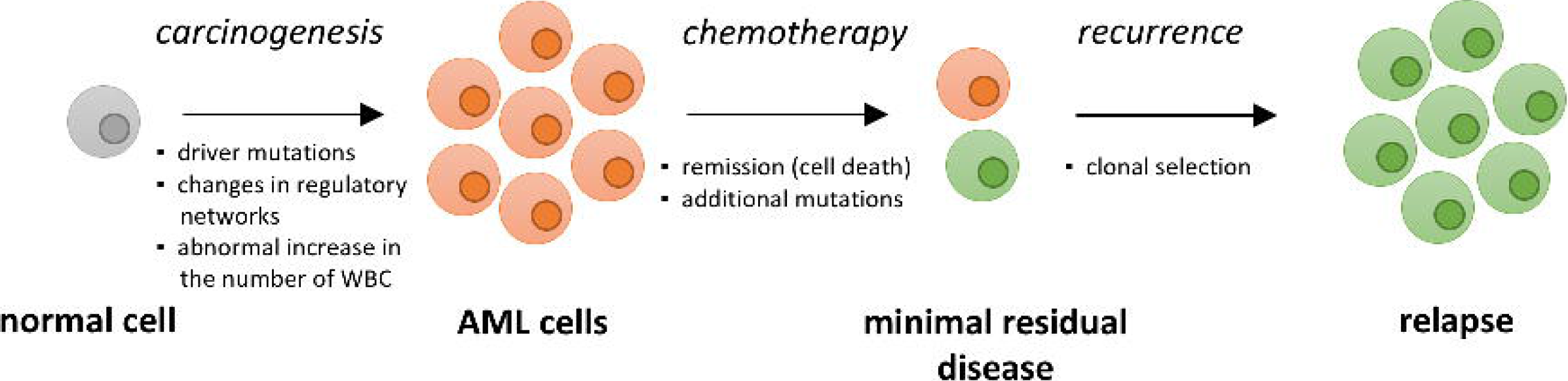
Sequence of events in acute myeloid leukemia, including carcinogenesis, diagnosis followed by chemotherapy, and relapse. Leukemic clones arise and compete against each other and healthy hematopoietic clones. Our mathematical model starts at diagnosis, so it does not include carcinogenesis. We also assume no additional clones arise by mutation after diagnosis.

Underlying our model are assumptions regarding the structure of growth, differentiation, and competition of the normal and leukemic clones. These are depicted in a schematic way in **Figure 2** (to be discussed in detail in **Methods**). Because human bone marrow cell counts are too large for direct stochastic simulation methods to be effective, we developed a hybrid numerical algorithm combining stochastic Gillespie-type (12) and tau-leaping algorithms (13), and a deterministic differential equation solver, which uses much less computer time than a “straight Gillespie algorithm” (see **Methods** for more detail). Our model was fitted to somatic mutations data from patients enrolled in recent clinical trials available in The Cancer Genome Atlas (TCGA) and the Genotypes and Phenotypes (dbGaP) databases.

**Figure 2.**
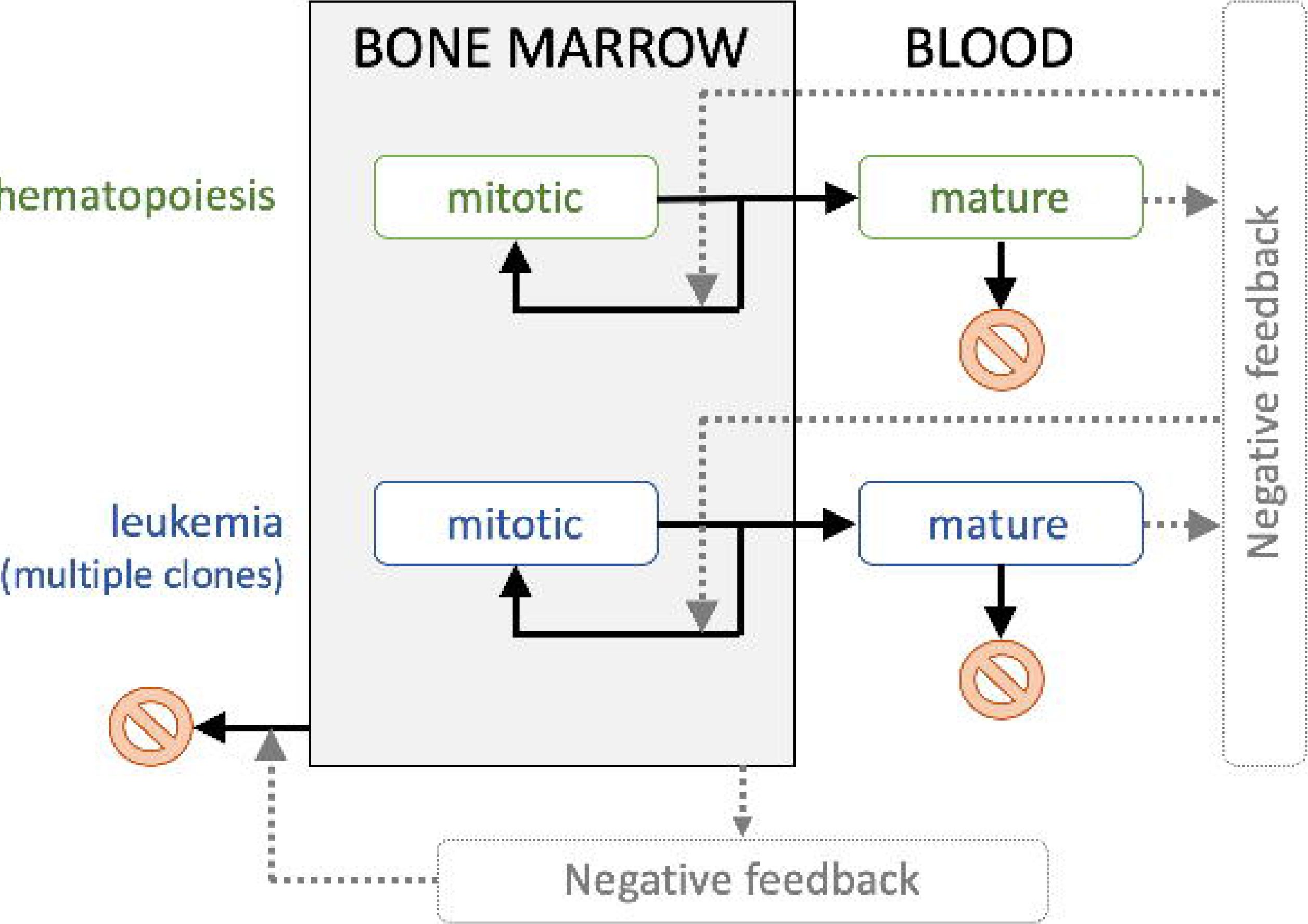
Schematic of the model. The negative feedback controlling the total population in blood downregulates all clones’ self-renewal rates if blood is overcrowded. The negative feedback controlling the total population in bone marrow upregulates the death rates of all compartments in bone marrow if it is overcrowded. Feedback configuration assumed follows closely (7).

Stochastic modeling can predict the range of outcomes of chemotherapy, given mutational data at diagnosis. The algorithm can be tested as a predictive tool for clinical management of patients with post-remission AML. To inform the model, we employ whole genome sequencing and clonal evolution data from the TCGA database. One limitation to existing models is the paucity of serial, paired sequencing data. Of 200 cases of AML sequenced at diagnosis, 20 patients were also sequenced at relapse, and only six have the clinical information required for the model available.

## Results

### The mathematical model can account for heterogeneity in clonal evolution

We develop a stochastic model for the evolution of leukemic clones along with the healthy hematopoietic one from diagnosis through treatment to relapse. Our model reflects the stochasticity inherent when leukemic clones are near depletion after chemotherapy, which we hypothesize strongly contributes to the interpatient heterogeneity in treatment response. The parameters are estimated by fitting the expected-value model (1–5) to the patient’s clinical data, as outlined in the **Methods**. Since the fits are not unique (see below), three sets of parameters for stochastic simulations are acquired from the list of expected-value fits, correspondingly characterized by low, average, and high renewal rates (see below for details).

### The input of the model includes clinical data and clonal landscapes at diagnosis and relapse

A set of clinical parameters (**Table 1**) was extracted from the TCGA database and used as input for modeling. The available data at diagnosis includes patient’s weight, percent cellularity, white blood cell count, percentage of blasts in both peripheral blood and bone marrow, and percentage of normal neutrophils in the peripheral blood. Importantly, the time to relapse and percentage of blasts in bone marrow at relapse are available. **Table 2** lists each patient’s parameters estimated based on the data in **Table 1**. These serve as input parameters for the stochastic model and the fitting procedure. Several constants for estimating the cell populations at diagnosis per kilogram of body weight are obtained from (7) and adjusted here using the patient’s weight from the TCGA dataset.

**Table 2:** Parameters computed for each patient based on TCGA data.

| Parameter | Calculation | Interpretation |
| --- | --- | --- |
| $\bar{c}_1$ | $= 2 \times 10^9 \times w$ | equilibrium normal mitotic cell population in the absence of leukemia<sup>a</sup> |
| $p_{diag}^{M,total}$ | $= 4.6 \times 10^{11} \times \%_{cellularity}$ | total cell population in BM at diagnosis<sup>b</sup> |
| $p_{diag}^{B,WBC}$ | $= 5 \times 10^9 \times WBC$ | total population of WBC in blood at diagnosis<sup>c</sup> |
| $p_{diag}^{B,blast}$ | $= p_{diag}^{B,WBC} \times \%_{blast}/100$ | total population of blasts in blood at diagnosis |
| $p_{diag}^{B,normal}$ | $= p_{diag}^{B,WBC} \times \%_{segs/neutrophils}/100$ | total population of normal cells in blood at diagnosis |

<sup>a,b</sup>The values arise from calculation of the hematopoietic cell lineage per kilogram of body weight in (7).
<sup>c</sup>Based on the assumption that, on the average, the patients have 5l of blood.

To simplify parameter estimation, we employed expected-value model trajectories that satisfy a system of ordinary differential equations (ODEs). As detailed in **Methods**, we first identify leukemic clones by clustering in the variant allele frequency (VAF) space using the Mclust algorithm. For each clone, we estimate two parameters: the proliferation rate *p*^*l*^, at which the clone’s mitotic cells divide, and the self-renewal rate *a*^*l*^, which is the probability of each daughter cell becoming a new mitotic cell as opposed to becoming a mature cell. Estimates of these parameters, specified and generally different for each leukemia clone, are obtained by fitting the model to data. The death rates for all leukemic clones are assumed to be equal (see **Methods**). This assumption is unlikely to hold true for actual leukemic cell populations, but it helps minimize the number of parameters for fitting per patient. Finally, the fit must satisfy the biological constraint of remission, which all patients achieved after the consolidation period. This requires that the total leukemic population at the end of treatment constitute less than 5% of the total bone marrow mononuclear cell population, corresponding to a classical morphologic definition of remission.

Uncertainty of fitting is caused by the fact that different estimates of self-renewal and proliferation rates for the leukemic clones can lead to the same results at relapse. In each case, the fit obtained is of desired quality, as described in **Methods**. As shown further on, the configuration of estimated pairs (*a*^*l*^, *p*^*l*^) depends on the number of clones present at diagnosis and relapse. The uncertainty is caused by an inability to distinguish, using data available, the two following extreme scenarios (as well as a range of intermediate ones):

**Scenario 1:** Clones that exist at diagnosis but not at relapse are those that are not competitive compared to the other clones. Generally, the corresponding fits include low renewal and proliferation rates.
**Scenario 2:** Clones that exist at diagnosis are not present at relapse because they are the most competitive, i.e. they have renewal rates or higher proliferation rates higher than the other leukemic clones. Therefore, under chemotherapy, with the cell kill assumed proportional to each clone’s proliferation rate (Eqs (9–10)), the mitotic population modeled is decreasing sharply and hence it is eliminated.

Uncertainty of parameter fits affects the distribution of outcomes of the stochastic model, while it does not affect the fit quality, although it generally affects the expected trajectories at times between diagnosis and relapse. These effects are separate from the intrinsic randomness of the stochastic simulations starting from MRD levels. In further analysis of the stochastic results, we choose 3 parameter sets from the 100 found using this parameter fitting process. The chosen parameter sets can lead to qualitatively different outcomes for the patients.

### Estimation of coefficients

For each patient, 100 parameter sets are found using the parameter fitting scheme. All the parameters found for each patient are presented in **Figure 3**. There are several observations that can be drawn:

**Figure 3.**
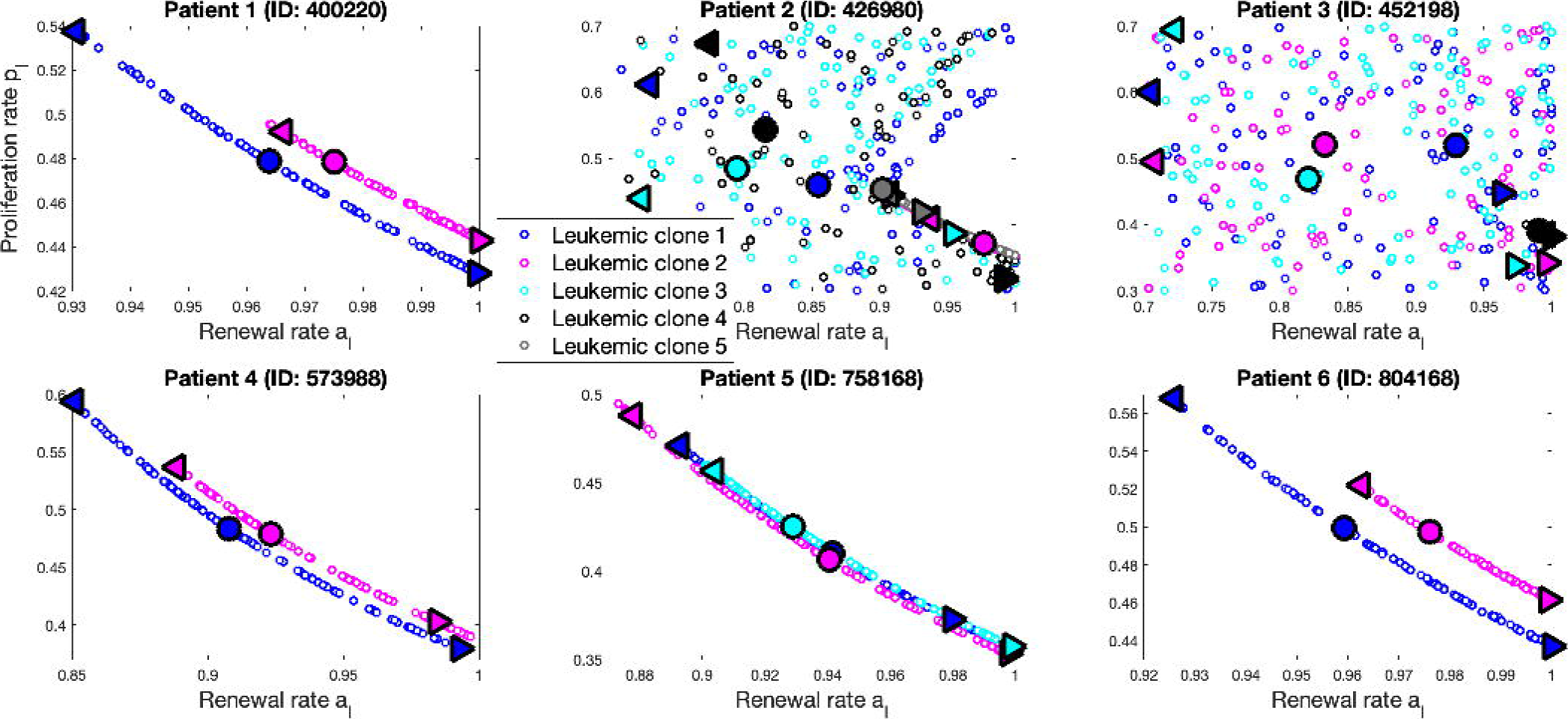
Parameter estimates for all patients. 100 parameter sets have been found for each patient starting from randomly selected initial conditions of the fitting procedure. Parameter sets corresponding to clones present at both diagnosis and relapse form curves in the parameter space, while those corresponding to clones absent at relapse display a more complicated pattern, implying different scenarios for these clones to become extinct. For each patient, three parameter sets have been chosen for stochastic simulations, corresponding to low, medium and high renewal rates (left triangles, circles, and right triangles).

For the clones which are present at relapse, the pairs (*a*^*l*^, *p*^*l*^) form curves in the parameter space. This phenomenon was also observed in (14). A given clone curve is above others if the clone is more competitive (higher renewal rates or higher proliferation rates; see Patients 400220, 573988, and 804168 in **Figure 3**).

For the clones which, in the model, become extinct at relapse, there is no pattern to the pairs (*a*^*l*^, *p*^*l*^) (see Patients 426980 and 452198 in **Figure 3**). The pairs that are either above the curves or slightly below the curves correspond to Scenario 2, in which the corresponding clones are extremely competitive and die during chemotherapy because of the high proliferation rates. The pairs that are well below the curves correspond to Scenario 1, in which the corresponding clones die because they cannot compete against the other leukemic clones or the normal clone.

**Figure 4** shows one expected trajectory for Patient 3 corresponding to a particular parameter set. For the same patient, different parameter sets result in different dynamics in the expected trajectories, however they all have the same clonal percentages at diagnosis and relapse, since this is the goal of the fitting scheme. Under this parameter set, the model predicts that leukemic clone 1 is eliminated during treatment, clone 2 escapes treatment but grows very slowly and is undetectable at relapse, clone 3 also escapes treatment but was subsequently outcompeted by the other clones, and clone 4 grows steadily from a small level after remission and is the only leukemic clone detected at relapse.

**Figure 4.**
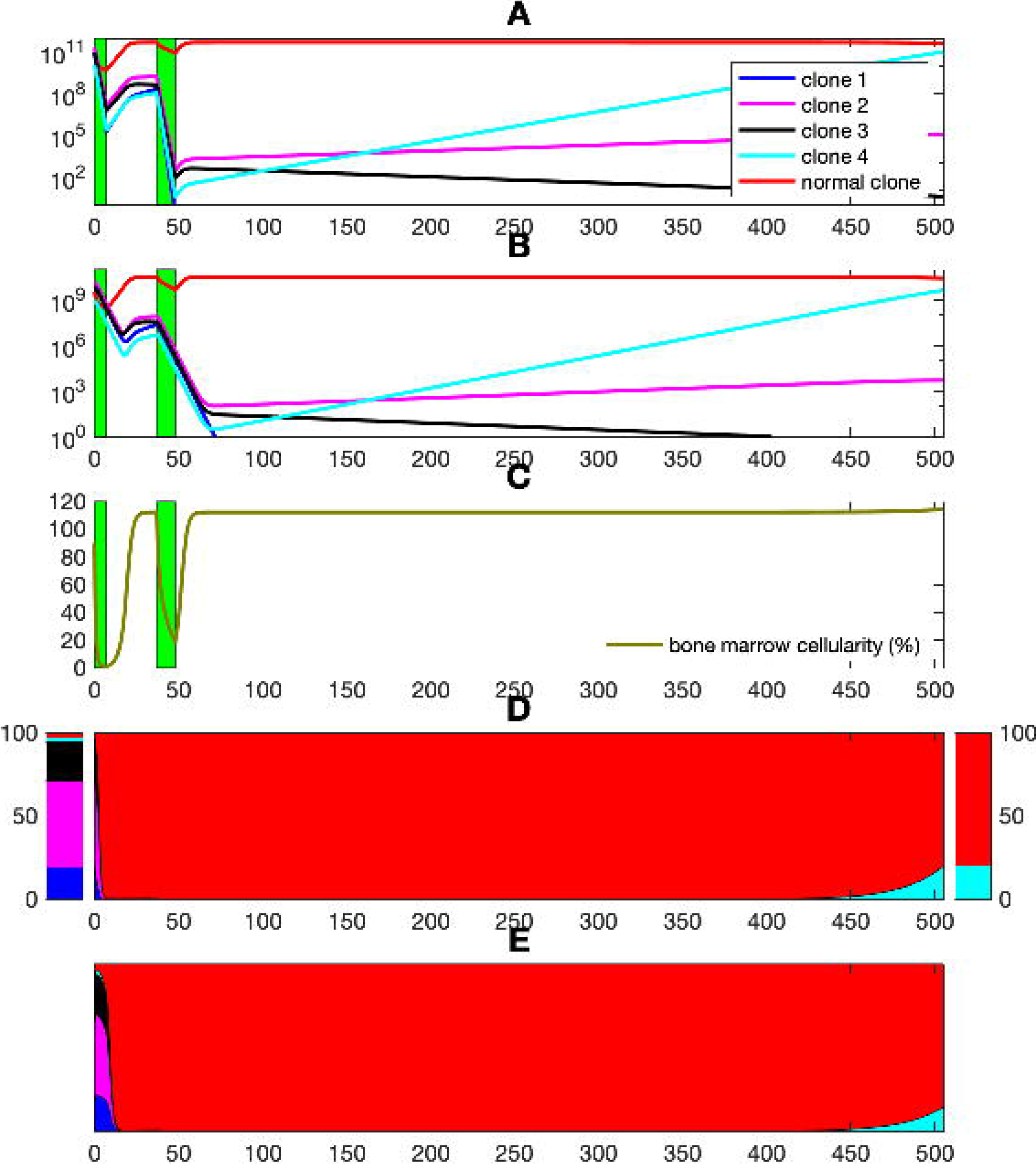
Results of the fitting procedure for Patient 3 (ID: 452198). Example of fitting the expected trajectories using a single parameter set. **(A)** Evolution of mitotic populations in the BM of all clones, with cell counts in logarithmic scale. **(B)** Evolution of the mature populations in peripheral blood. Green bars indicate chemotherapy treatments. **(C)** Evolution of BM cellularity. **(D)** Evolution of clonal percentages in bone marrow. The bar-plot on the left consists of clonal percentages at diagnosis, and the one on the right consists of clonal percentages at relapse (both are based on sequencing data). The parameter sets are chosen to fit the clonality data in these two bar-plots, as shown in the middle plot. The color code for different clones in (B) and (D) is the same as in (A), as described in its legend. **(E)** Evolution of clonal percentages in peripheral blood.

### Stochastic simulations

As detailed in previous sections, for each patient 100 parameter sets were found. Three of the parameter sets are chosen such that leukemic clones typically have

Parameter set 1: high proliferation rates and low renewal rates
Parameter set 2: intermediate proliferation and renewal rates
Parameter set 3: low proliferation rates and high renewal rates

For each parameter set, 1000 stochastic trajectories have been created by using the hybrid stochastic simulation algorithm, described in **Methods**. These trajectories are different paths of remission achieved by chemotherapy, nevertheless with the disease progressing to relapse. The trajectories are categorized into disjoint “outcomes”, based on the number of mitotic cells in each clone at relapse. For demonstration purposes, a clone is considered dead at relapse if it has less than 1000 mitotic cells, and alive at relapse otherwise. Using a different threshold does not lead to a marked change in the results.

**Figure 5** shows the expected-value and stochastic outcomes for Patient 5. Under parameter set 1, leukemic clones have high proliferation rates and low renewal rates. The result is that the clones are highly affected by chemotherapy (since mitotic cell death is assumed to be proportional to proliferation rate during treatment). In the expected-value model, the disease is almost eradicated but still exists in a very small population at remission, which eventually gives rise to relapse. As a result, no stochastic simulation leads to the outcome where all leukemic clones detected at diagnosis are present at relapse, which is predicted by the expected-value model. Because the simulated leukemic population is small at remission, random fluctuations lead to eradication of one or more leukemic clones soon after treatment. For parameter set 3, in which clones have low proliferation rates and high renewal rates, chemotherapy has smaller impact on the leukemic populations. The disease exists in larger percentages at remission, and gradually increases in size until relapse. Parameter set 2 presents a middle ground between parameter sets 1 and 3: the disease is almost eradicated by chemotherapy, such as in parameter set 1, but remission contains a larger leukemic population and therefore all clones are more likely to progress to relapse. 64.1% of stochastic simulations lead to the same outcome as the expected-value simulation.

**Figure 5.**
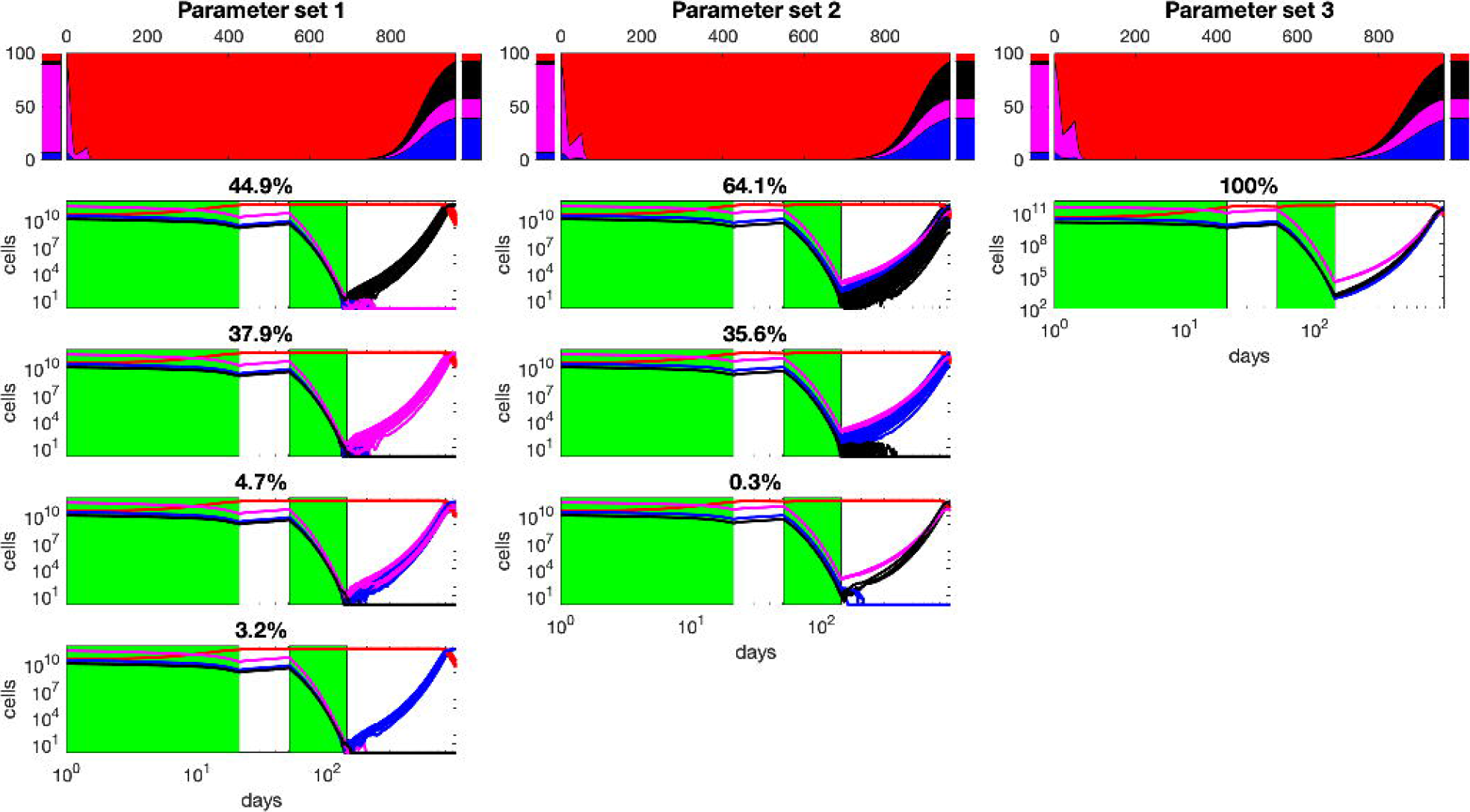
Results of the stochastic model for Patient 5 (ID: 758168). Columns correspond to different parameter sets. Row 1: Evolution of clonal percentages in the expected-value model. Rows 2 and on: Outcomes of the stochastic model, with cell counts in logarithmic scale, with corresponding frequencies of occurrence listed. Stochastic simulations agree with the expected-value results in parameter set 3. In parameter set 1, no simulation leads to the same outcome as the expected-value simulation, where all three leukemic clones detected at diagnosis are present at relapse. This increases to 64.1% in parameter set 2 and 100% in parameter set 3. The parameter sets for all patients are listed in Supplemental Table S1.

**Figure 6** shows the expected and stochastic outcomes for Patient 2. The expectation results show that only clones 2 and 5 are present at relapse. 42.4%, 92.3% and 0% of the stochastic simulations under parameter set 1, 2 and 3 lead to this outcome, respectively. Under parameter set 1, all other leukemic clones are eradicated by chemotherapy. However, random fluctuations also may lead to either clone 2 or 5 being eliminated even after treatment. The stochastic simulations under parameter set 3 follow a different route. No leukemic clone is entirely eradicated by the time of complete remission, but clones 1 and 4 are gradually outcompeted. Furthermore, clone 3 is present at relapse in all stochastic simulations, but at population sizes smaller than the detection level. Similarly as for parameter set 1, clones 2 and 5 can each be erased due to stochastic fluctuations.

**Figure 6.**
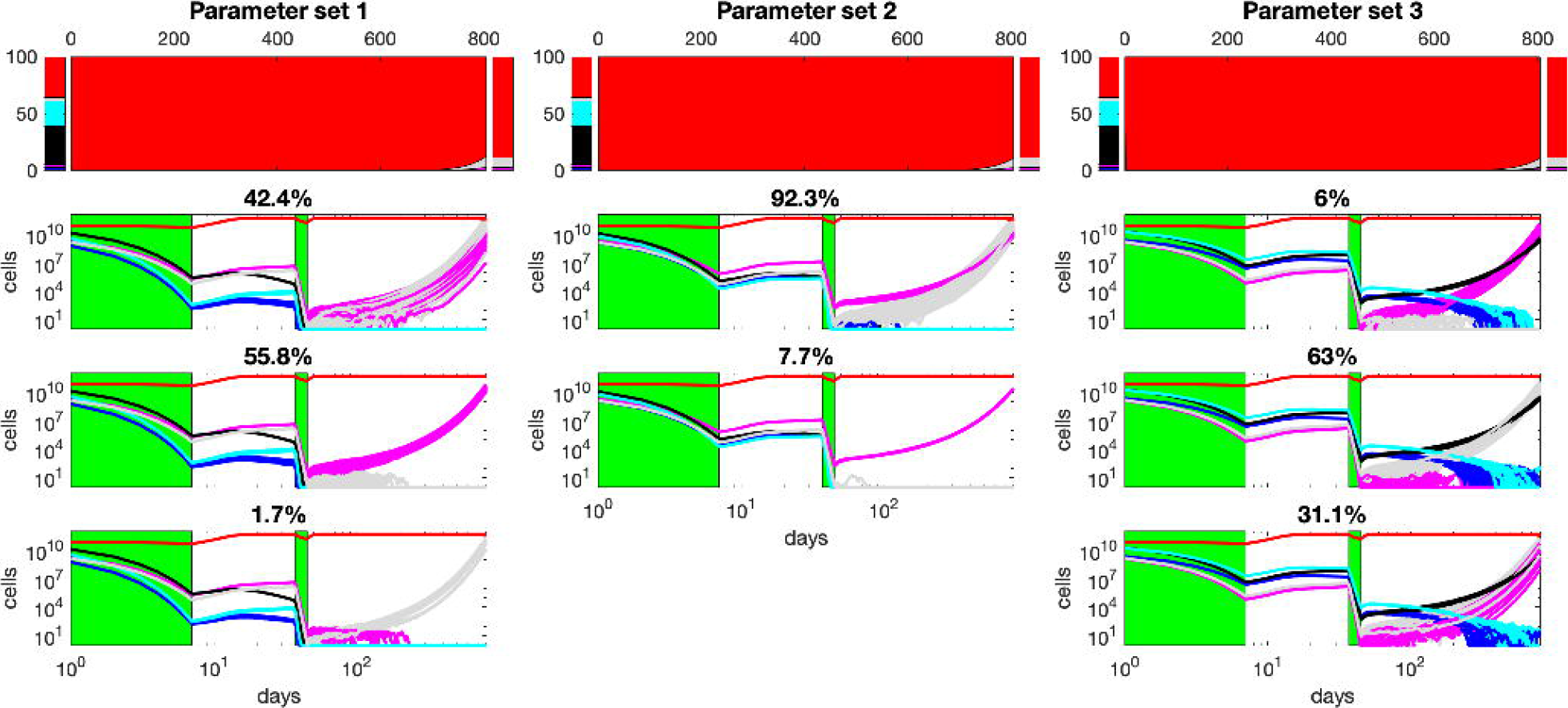
Results of the stochastic model for Patient 2 (ID: 426980). Columns correspond to different parameter sets. Row 1: Evolution of clonal percentages in the expected-value model. Rows 2 and on: Outcomes of the stochastic model, in logarithmic scale with corresponding frequencies of occurrence listed.

### Comparison to independent estimates of the MRD and clone proliferation rates

Ivey et al. (15) studied patient bone marrow at several time points following remission to detect MRD as early as possible. All patients had AML with mutated *NPM1* gene. This study allowed drafting approximate trajectories leading to hematological relapse. Under assumption that the re-growth of the malignant clone is exponential, the resulting growth rates are included in the range of 0.3 to 2.0 log_10_ month^−1^, as reproduced in **Figure 7.**

**Figure 7.**
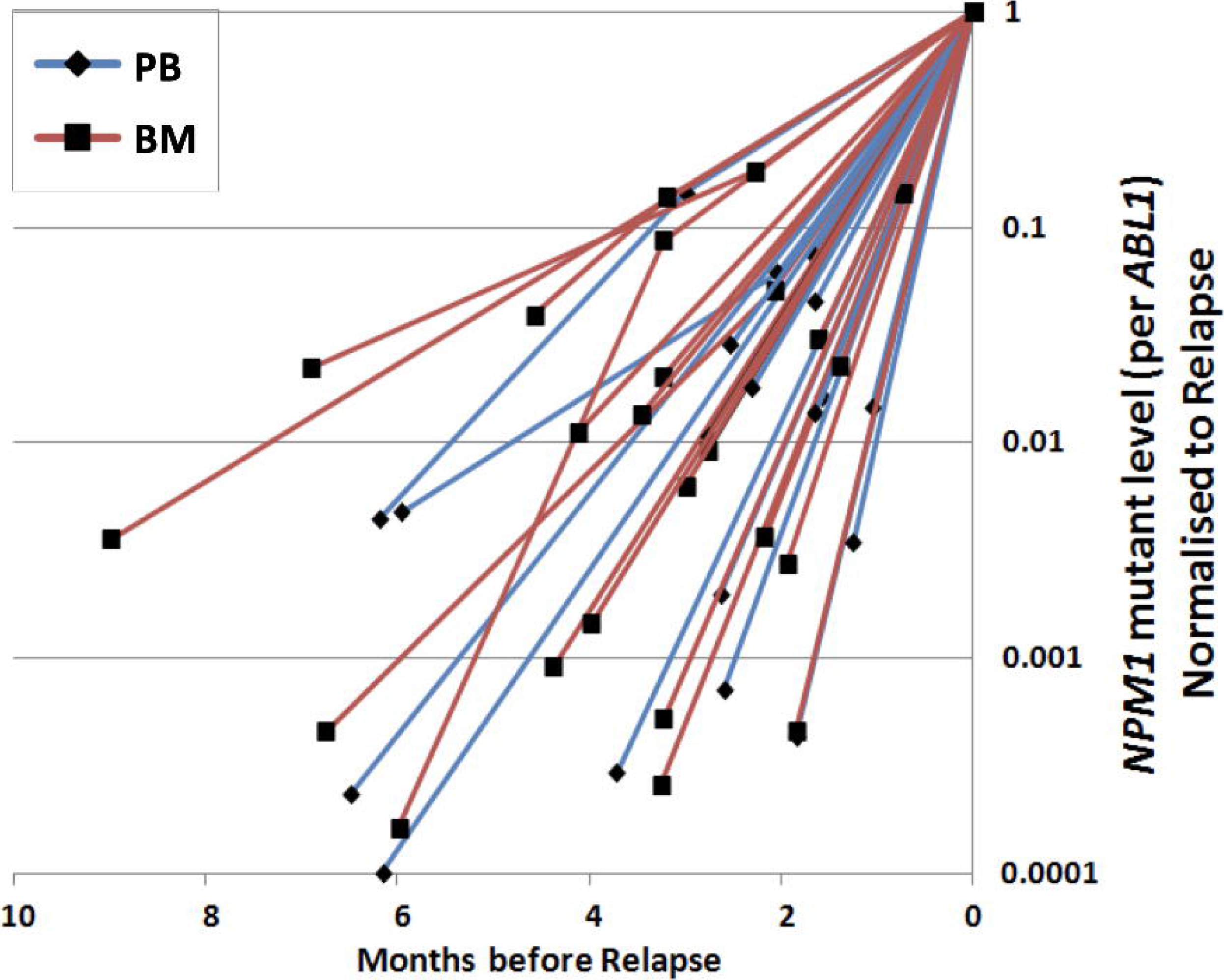
Kinetics of relapse of NPM1-mutated AML (reproduced from (15) with permission). Based on sequential monitoring of samples obtained in human patients after the end of chemotherapy until molecular or hematological relapse. PB, peripheral blood; BM, bone marrow.

We compared the growth rates of the clones in the TCGA AML patients to see if they fit into this range (assuming clones with different mutations have similar growth rates to the *NPM1*-mutant clones). Only the clones still present in the patients’ clinical data at relapse were considered. Their growth rates can be computed from the parameters resulting from model fitting (see **Methods**).

When compared with the result from Ivey et al. (15), the growth rates for all patients in our analysis fit in the range of experimentally observed data (**Figure 8**). Growth rates are smaller when it takes longer for the disease to relapse in a given patient, and vice versa.

**Figure 8.**
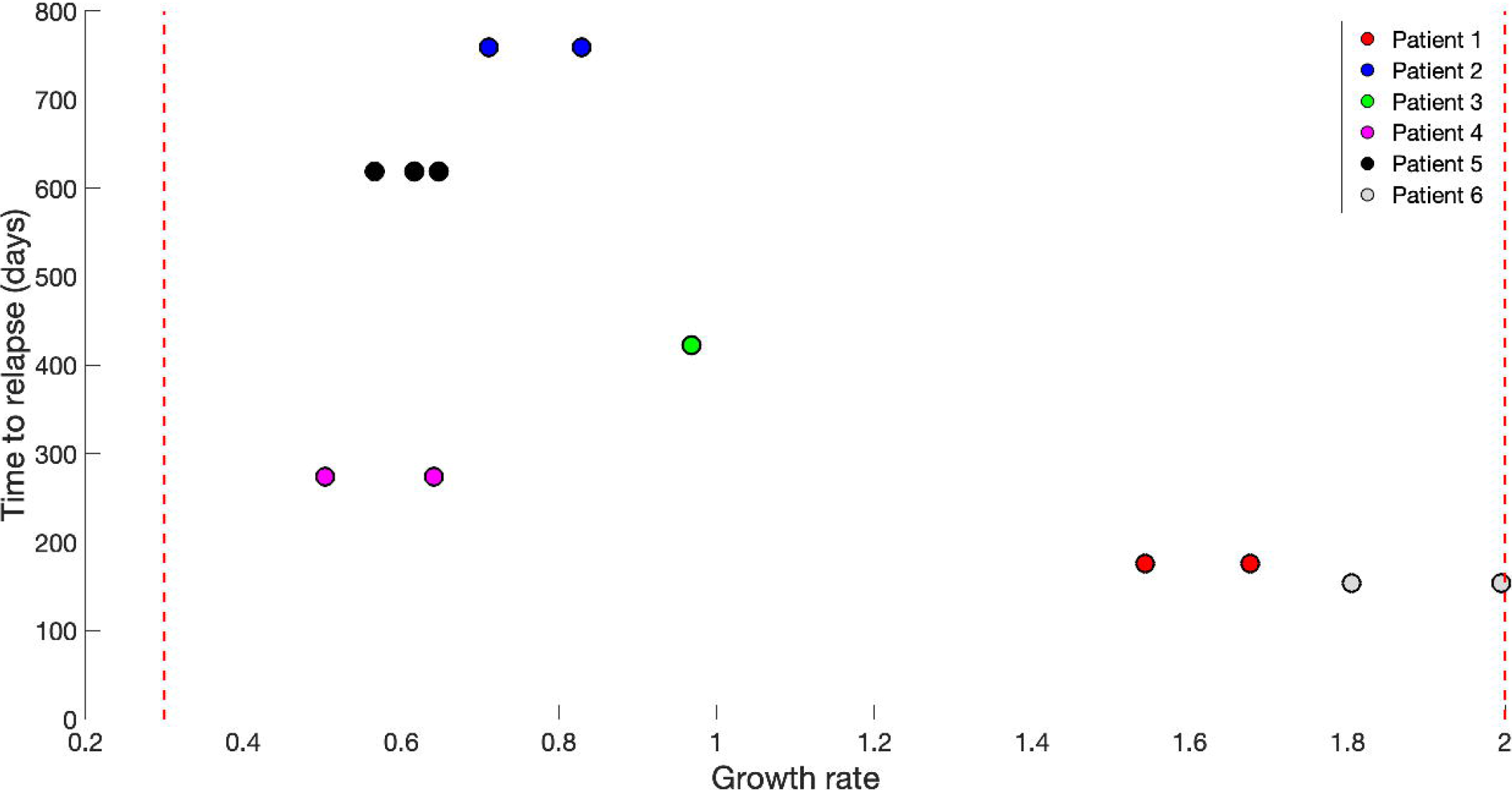
Inferred growth rates of different clones in all patients. Only the clones that still exist at relapse in the patients’ data were considered. The red lines are bounds for growth rates from 0.3 to 2.0 log_10_ month^-1^, based on the range of growth rates of NPM1 mutant clones in (15); see Fig 7. Growth rates computed from our model (blue circles) fit into this range.

Computed growth rates agree with experimental data, which is an indication that the parameters we derived from the expected-value model might be biologically relevant.

### Relationship between MRD and the time to relapse

We next study the connection between MRD (defined as the clonal percentage in BM at 6 months after remission) and time to relapse (defined as the time period from remission to when %BM blast exceeds 5%) inferred from the stochastic model (3 parameter sets per patient, as explained earlier on). **Figure 9** shows the comparison in all patients.

**Figure 9.**
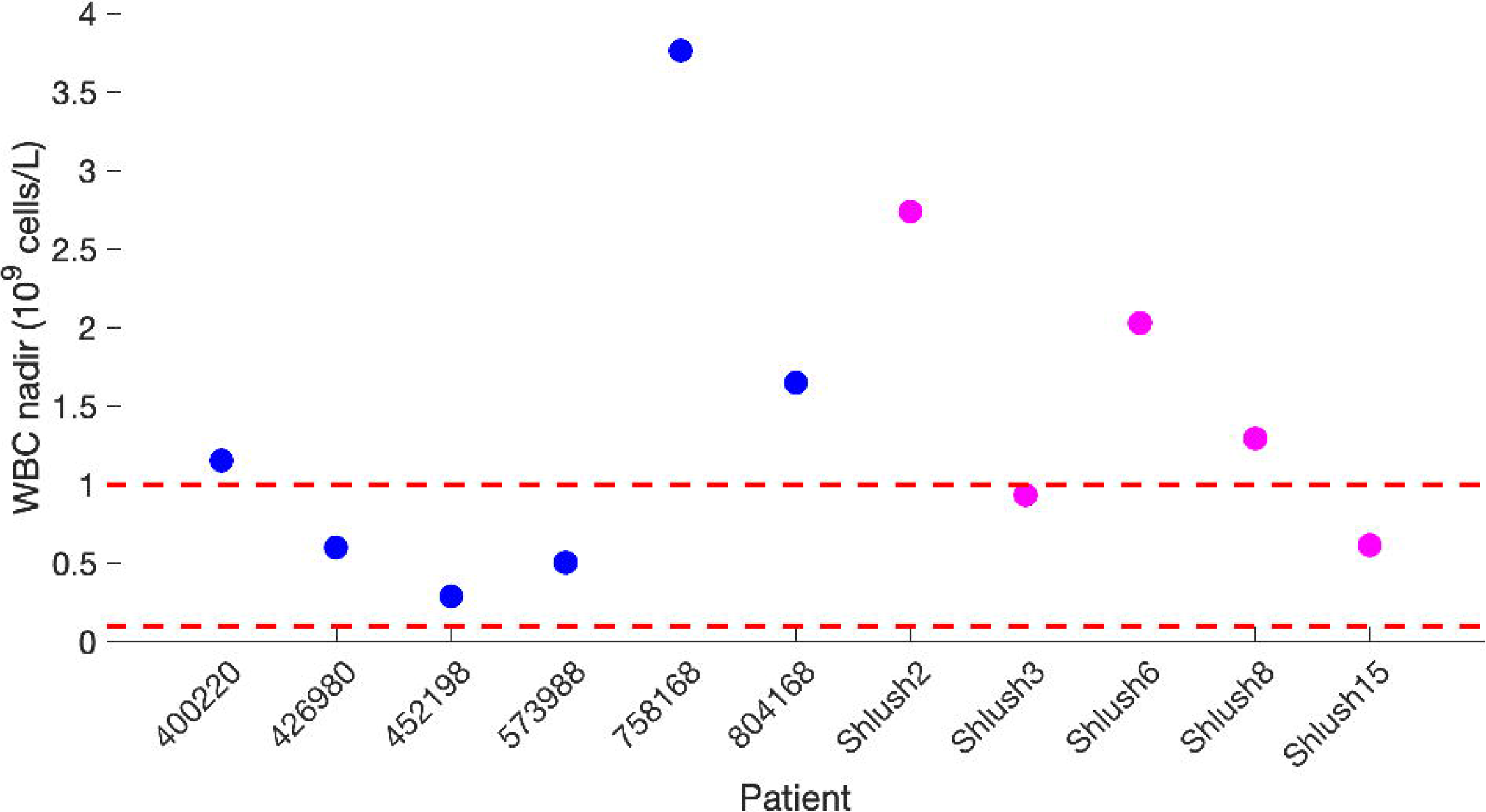
Relationship between MRD and time to relapse for all patients. Estimates of MRD and time to relapse are mean values from 1000 stochastic simulations. Each color corresponds to a single patient; left triangles, circles, and right triangles correspond to parameter sets 1, 2 and 3, respectively. Simulated points are fitted with a sigmoidal Hill function of the form: Time to relapse = *A*/(*B* + (log(*MRD*) − *C*)^*n*^) where *A* = 15539, *B* = 20, *C* = −7 *n* = 2.

For all patients, the MRD at 6 months is smallest in parameter set 1 and largest in parameter set 3. This is in agreement with our previous observation from the stochastic model; parameter set 1 results in leukemic clones being affected the most by chemotherapy and therefore the disease level at remission is very small (some clonal populations can decrease to ~10^1^ − 10^2^ cells; see Figs. 4 and 5). Comparing all patients, it is clear that larger MRD is associated with shorter time to relapse.

### Analysis of MRD in bone marrow and peripheral blood

Finally, we analyze the blast percentage in both bone marrow (%BM blast) and in blood (%PB blast). **Figure 10A** shows the evolution of %BM blast and %PB blast in Patient 3 (ID: 452198). Also shown here is the frequency of %BM blast when % peripheral blood (PB) blast reaches 5% in the stochastic results, usually defined to be the time of relapse (**Figure 10B**).

**Figure 10.**
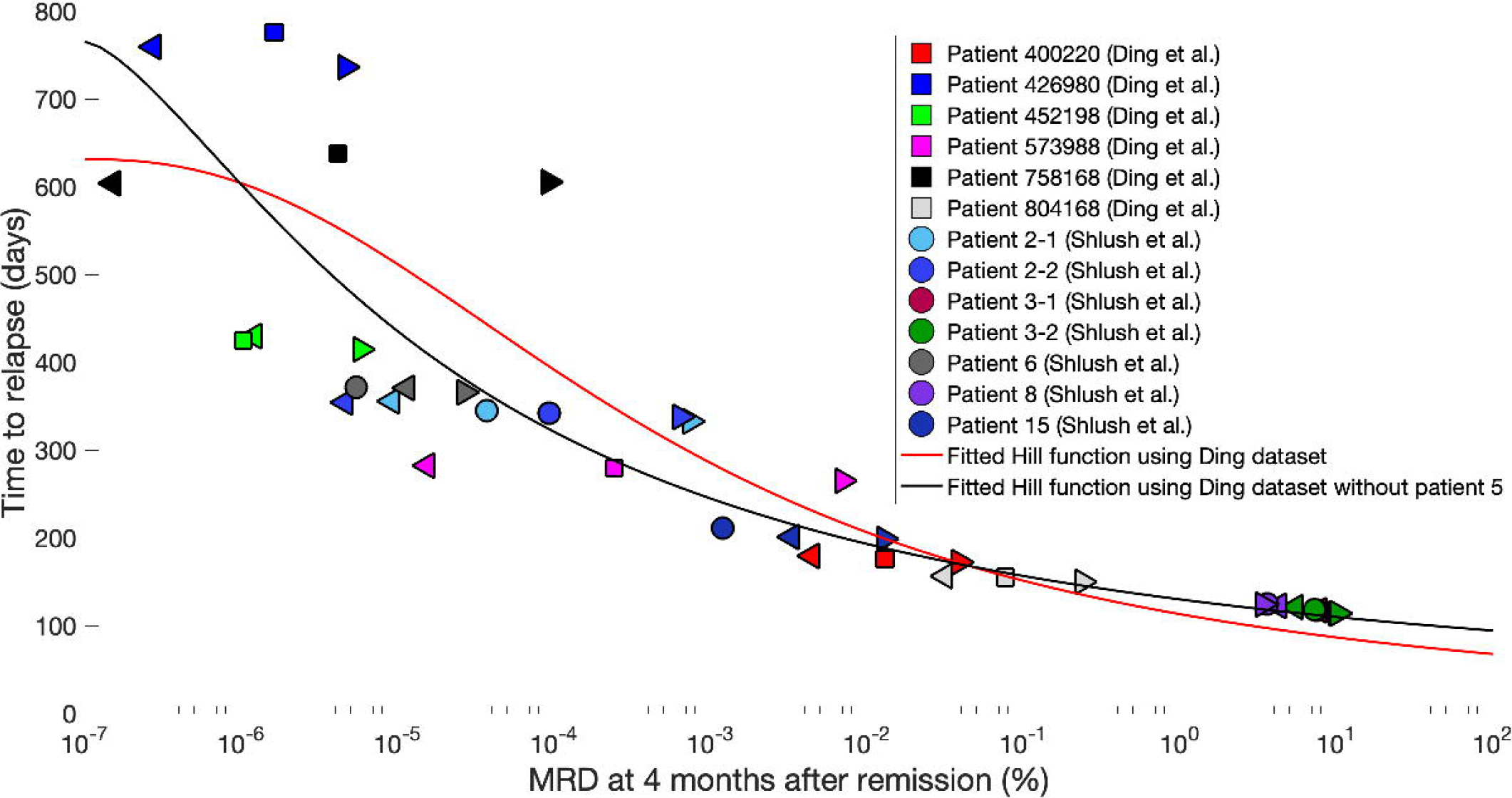
Analysis of MRD for Patient 3 (ID: 452198). ***(A)*:** Evolution of %BM blast (red) and %PB blast (blue) from diagnosis to relapse. Solid lines: Mean values, broken lines: minimum/maximum values. The mean %BM blast and the mean %PB blast are largely close from complete remission to relapse. The minimum %BM blast and %PB blast indicate that some simulations lead to the disease being eradicated. ***(B)*:** Frequency graph of %BM blast when %PB blast reaches 5%.

In Patient 3, a number of stochastic simulations lead to eradication of some leukemic clones. Because of this, the minimum %BM blast and %PB blast are effectively at 0% after chemotherapy. In spite of this, the mean %BM blast from simulations leads to the same value as that observed in clinical data. The mean %BM blast and the mean %PB blast are largely identical from the end of chemotherapy to relapse. As a result, when %PB blast reaches 5% in the stochastic simulations, the %BM blast is also distributed close to 5% (**Figure 10B**).

## Discussion

Despite major advances in leukemia genetics and molecular targeted therapies, event-free and overall survival of AML patients remains less than 40%. Emergence of chemoresistant clones and relapse after intensive chemotherapy or stem cell transplantation remain the chief obstacles. Identification of MRD as an independent prognostic factor, stratification of patients by their status, and early intervention with change in therapy has improved outcomes in acute lymphoblastic leukemia (16–22). A challenge has been how to define MRD in AML. The threshold of what constitutes MRD in AML is not well-established in the terms of either cytogenetics, or leukemia-associated immunophenotype, or mutational allelic frequency (20). Each of these assays has a different level of sensitivity. Current clinically-accepted definition of MRD at 0.01% is based on flow cytometric leukemia-associated immunophenotypes. Different subtypes of AML, e.g., *NPM1* or *FLT3* mutations, may have different MRD dynamics. Relapse can be associated with loss of founder mutations (23). Somewhat surprisingly, flow cytometric false-negative results occur in almost 19% (19). Nonetheless, a growing number of clinical trials document the prognostic value of MRD in AML (20). One of our results is a model-based MRD – time to relapse relationship (**Fig. 9**), indicating large uncertainty in predicting the time to relapse. Making this prediction more accurate will require further data on clonal evolution in bone marrow from diagnosis to relapse.

Here, we report our use of patient-derived data to design stochastic modeling based on age-dependent Markov branching process. The structure of our model is similar to that by Stiehl et al. (7), which can be considered a most parsimonious model taking into account treatment and competition of normal and leukemic clones in the bone marrow. However, our model is entirely stochastic. The data we employed to estimate our model parameters include treatment details, blood counts, and bone marrow biopsy results from which the clonal landscapes at diagnosis and relapse can be determined. The model was calibrated to fit the clonal frequencies, while satisfying the biological requirements as observed in the data. This allows us to realistically reconstruct the rise and fall of clonal populations, and take into account uncertainties of estimation process, as well as those inherent in AML relapse having a random component.

The parameters fitted to the expected-value model offer an explanation of how a leukemic clone can escape chemotherapy and promote relapse. As it has been observed in (14), these clones either have high proliferation rates or high self-renewal rates. As a result, there is a range of different parameter combinations that can explain their ability to succeed. On the other hand, we also study the clones that have been eradicated by the time of relapse and conclude that these clones might be eliminated either because they are not competitive and therefore surrender to other clones, or they are simply killed by chemotherapy. Also, we checked if the parameters are biologically relevant by using the model to compute the corresponding clonal growth rates for each patient. That these values fit in the clinically observed range independently found in (15) for patients with the *NPM1* mutations, suggests that the model is consistent with clinical data.

In (21), the authors modeled the emergence of clonal heterogeneity by modifying the ODE model to accommodate new leukemic clones arising from mutations. We do not include mutations in our model here because relapse more likely results from MRD than new mutations during and after chemotherapy (24, 25). The deterministic ODE models in (7, 14, 21) were used to track the evolution of different leukemic clones from diagnosis to relapse. This is an appropriate approach when the populations advance toward equilibrium. However, during and after chemotherapy, leukemic populations are experiencing bottlenecks and therefore are subject to stochastic fluctuations. Because of this, a stochastic branching process model was used, such that the expectations (means) of its trajectories approximately satisfy the ODE model. We showed that with the same parameters, the stochastic model predicts a variety of outcomes for the disease, due to the stochastic fluctuations in the MRD. This partially explains the inter-individual heterogeneity in response to treatment of AML and underlines the importance of a careful MRD monitoring in predicting the progress of the disease.

An interesting general conclusion from this analysis is once more that the “generalized” growth rate is not sufficient in understanding the effects of chemotherapy. It is necessary to distinguish between the division rate and self-renewal fraction. Various combinations of these two parameters lead to the same growth rates, but different treatment effects. This seems to be not universally recognized, although a review paper has been recently devoted to this important issue (26).

## Methods

### Data

Somatic mutation data for primary and relapsed tumor samples were derived from dbGaP (phs000159) for 8 patients, with the following IDs: 400220, 426980, 452198, 573988, 758168, 804168, 869586, 933124. Clinical information, including treatment details, complete blood count and bone marrow biopsy, was obtained from dbGaP and TCGA and supplements to (22, 27) and summarized in **Figure 11**. Tumor clone frequencies were calculated from deep read count data in (22). The somatic mutation variant allele frequencies (VAF) at primary and relapsed tumors are clustered by the Gaussian mixture based Mclust algorithm (available from the CRAN repository of R-codes under https://cran.r-project.org/web/packages/mclust; also read the Supplement to (22) for more details). Patients 869586 and 933124 were excluded because they underwent autologous hematopoietic stem cell rescue after chemotherapy but before relapse, so they do not seem directly comparable with the other 6 patients.

**Figure 11.**
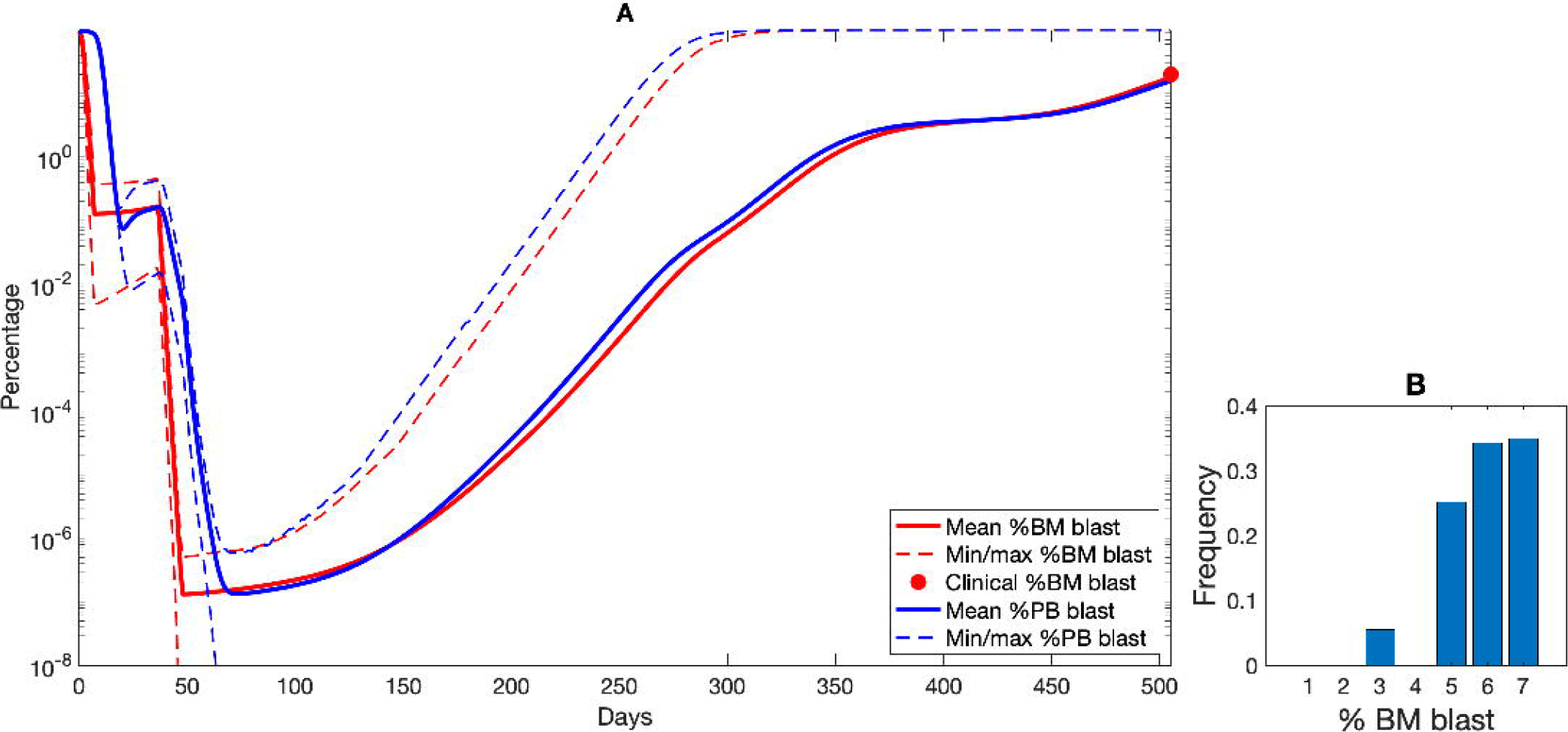
Detailed information about patients used in the study. Clinical information of individual patients. *Left:* Information about weight, age, sex, percentages of blasts at diagnosis and relapse, cellularity of bone marrow and percentage of segs or neutrophils. *Right:* Number of months since initial diagnosis to relapse and death of each patient, except for Patient 452198, whose history ends in 2011 with the last follow-up at which the patient was alive.

In the clonal analysis, some clones might be nested in others, because the former originate from the latter after gaining additional mutations. Since we define the clones as disjoint, the percentage of a clone at either time point is defined as the percentage of this clone with all subclones excluded. Therefore, the mutant clones’ and the normal clone’s frequencies at any given time point add up to 100%. An example of that is Patient 758168 (see **Figure 12B**) with three clones 1, 2 and 3. Clone 2 is a subclone of 1 and 3 is a subclone of 2. For this reason, clone 3 contains all mutations of clone 2 and 2 all of those in clone 1, however they show generally different allelic frequencies. To calculate the percentage of cells that belong to clone 1 we subtracted the allelic frequencies of clone 2 and 3 from clone 1 and multiplied them by 2 (assuming that most of the variants are heterozygous). Similar procedure was applied to all patients. Clone percentages were later scaled by the fraction of blasts in bone marrow, to additionally include the proportion of normal cells, since in all cases blast percentages were lower than 100%. We assumed that the relapsed clones do not include newly formed mutations, only that in the primary tumor some of them might have been undetectable. For this reason, cell percentages of clones with allele frequencies equal to zero were set to 3%.

**Figure 12.**
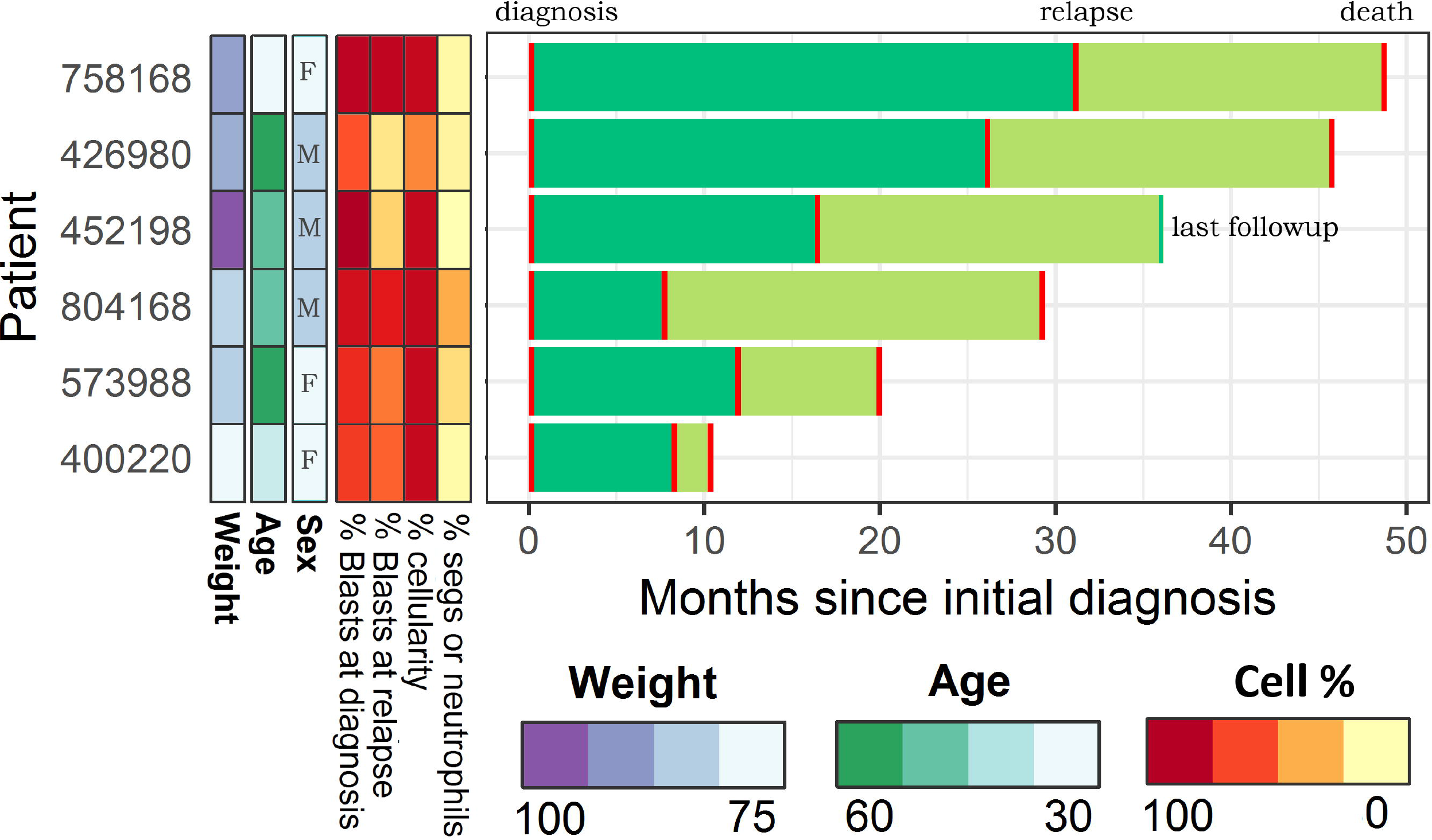
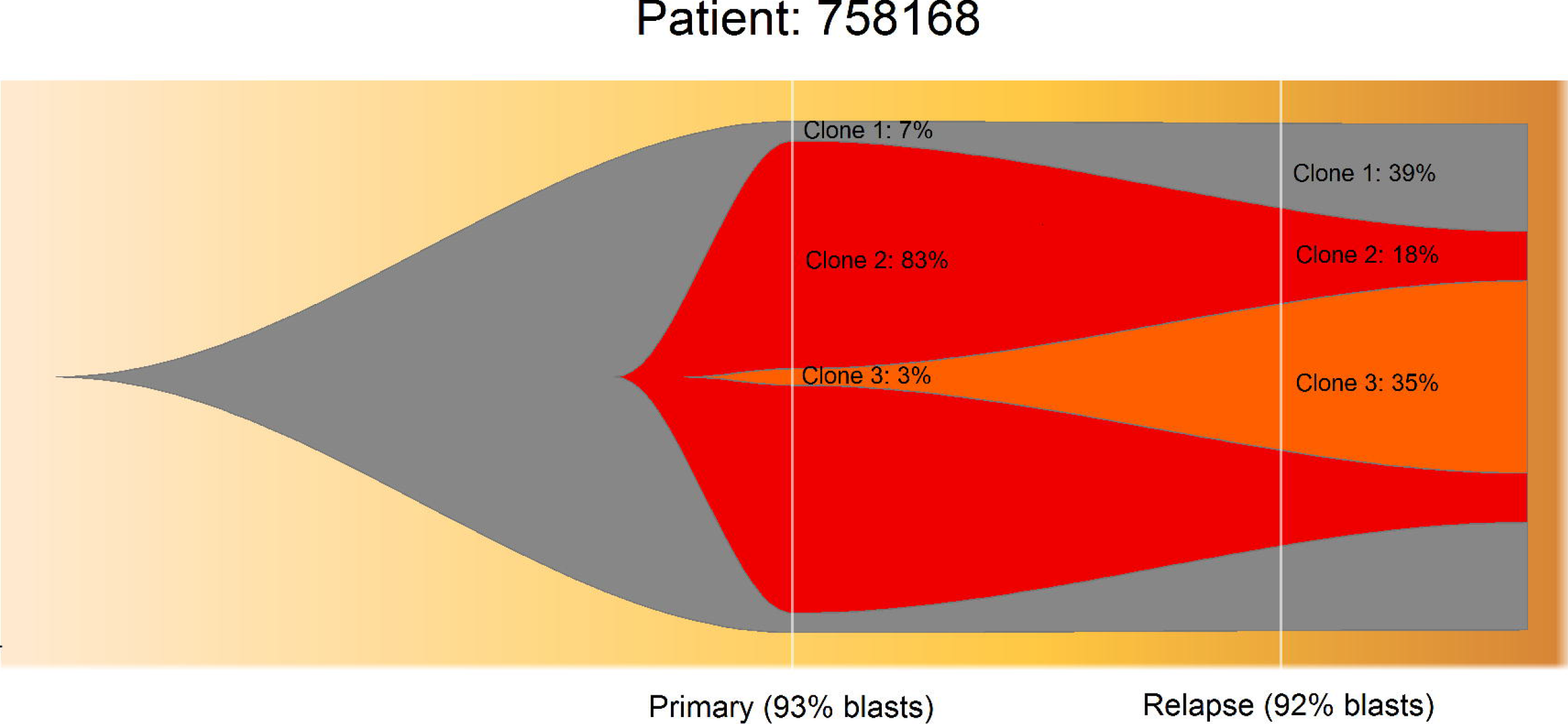
Clonal evolution for patient 5 (ID: 758168). Fractions of individual leukemic clones in primary tumor and relapse from whole-genome sequencing are represented as a fish plot, with percentage of each clone at either time point. Total width of the plot corresponds to the percentage of bone marrow blasts.

### Stochastic model for the proliferation and competition of normal and leukemic cells

The schematic of the stochastic model is depicted in **Figure 2**. Proliferation of cells in each clone in the model is represented by an ordered sequence of different compartments. Each event in the model (cell division, differentiation, or death) is characterized with an exponentially distributed waiting time, the rate of which will be explained below. Because the waiting times are random, the order in which reactions happen may differ between simulations. This results in a variety of outcomes under the same conditions, and may contribute to interpatient heterogeneity.

The two-compartment model for the hematopoietic clone was established in (28), in which the healthy clone is divided into two sub-populations, mitotic compartment *c*_1_(*t*) and mature compartment *c*_2_(*t*). The mitotic cell compartment, representing the more complex multi-stage differentiation process of hematopoietic stem cells (HSCs), hematopoietic progenitor cells (HPCs) and precursor cells, is located in the bone marrow (BM). Mitotic cells can divide into two daughter cells at the proliferation rate *p*^*c*^, and each of the daughter cells is either a new mitotic cell or a mature cell. The fraction of daughter cells returning to the mitotic cell compartment is called the self-renewal rate *a*^*c*^. The mature cell compartment consisting of neutrophil granulocytes, a major subtype of white blood cells, is located in the blood. Mature cells die at a constant rate *d*^*c*^. Each leukemic clone, as detected by sequencing data, also consists of two compartments: mitotic population *l*_1_(*t*) in BM, and mature population *l*_2_(*t*) in blood. The rules dictating its divisions, differentiations, and deaths, are similar to the hematopoietic clone, with proliferation rate *p*^*l*^, renewal rate *a*^*l*^and death rate *d*^*l*^.

There are two feedback systems governing the populations in blood and bone marrow. The first feedback system reacts to overcrowding in the blood by down-regulating the self-renewal rates of the hematopoietic and all leukemic clones by a factor of:

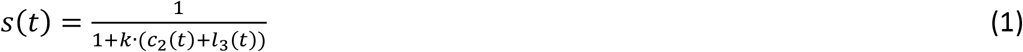

The second feedback system controls the total population in the bone marrow:

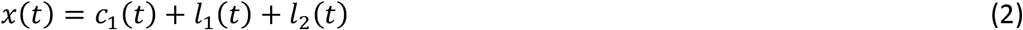

and if this population is too high, the death rates of all compartments in bone are increased by:

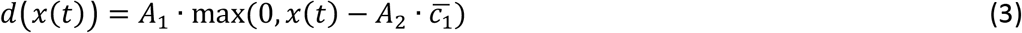

Finally, treatment drugs used in many chemotherapy protocols are characterized by increased killing of cells in the synthesis stage. Therefore, during treatment, the death rates of mitotic cells are increased by a factor proportional to their proliferation rates:

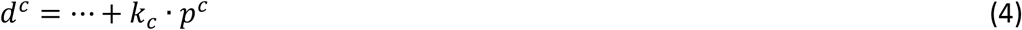

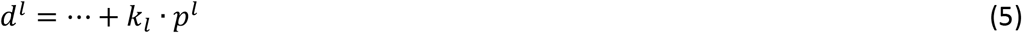

If any clonal mitotic population decreases below 1 cell during the time course, the clone is marked as dead and remains 0 until relapse. We assume that chemotherapy kills leukemic clones at a higher rate than the normal clone, therefore *k*_*l*_ > *k*_*c*_. Furthermore, we simplify the model by assuming a fixed ratio between the killing rates for all patients. The ratio *k*_*l*_: *k*_*c*_ = 5: 1 was found to result in realistic behavior of the disease trajectory.

### Expected-value approximation for fitting the stochastic model

While parameter estimation techniques have been extensively applied for deterministic models, such as those based on ordinary differential equations (ODEs), parameter fitting for stochastic models are still in early development. In this work, we approximate the stochastic model with a set of ODEs, use these expected-value approximations to fit the clinical data, then study the variability in the outcomes of the stochastic model under the resulting parameter sets.

The approximate expected-value dynamics of hematopoietic and leukemic compartments can be expressed from the division, differentiation and death events that govern their fluctuations, as described above:

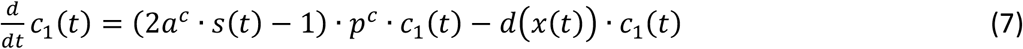

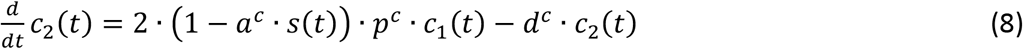

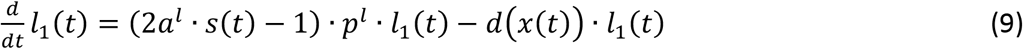

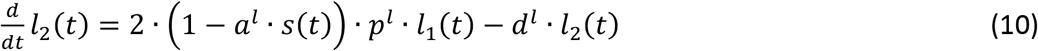

During chemotherapy, the mitotic populations further decrease:

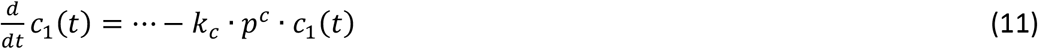

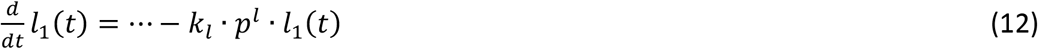

### Model parameters

1. Patient data from TCGA The data available in the TCGA dataset and used as inputs for our models are summarized in Table 1, including the differential counts in peripheral blood and lengths of chemotherapy treatments.
2. Dependent and free parameters of the model Stiehl et al. (7) calibrated the parameters for the hematopoietic cell lineage to data from the literature and concluded that for these cells, the self-renewal rate is *a*^*c*^ = 0.87, proliferation rate is *p*^*c*^ = 0.45 (*day*^−1^) and death rate is *d*^*c*^ = 2.3 (*day*^−1^). We use these parameters in our study.

The other important parameters for the model computed individually for each patient are listed in Table 2, with the equations to derive them from the TCGA dataset for each individual.

The feedback parameters are as follows:

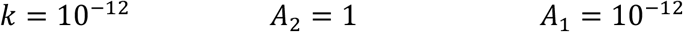

Different magnitudes of *k* were tested and we observed that the value *k* = 10^−l2^ produces simulations close to the patients’ clonal evolution. We chose *A*_2_ = 1 because if it is greater, the population in bone marrow can exceed its capacity. If *A*_1_ ≫ 10^−l2^, the log-plots of populations in bone marrow and blood develop sharp turns where they reach equilibrium values. In experimental data, it has been observed that these turns are more continuous, therefore we chose *A*_1_ = 10^−l2^.

The chemotherapy constants 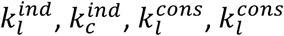, which model the effect of induction and consolidation treatments on the normal and leukemic cell lines, have been interactively determined depending on each patient’s clonal evolution. The free parameters of the model to be fitted for each patient are *a*^*l*^ and *p*^*l*^ of each leukemic clone. We observe that the magnitude of *d*^*l*^ may vary without noticeable consequences for the population evolution, because the fate of the disease depends on the leukemic mitotic population’s ability to grow in bone marrow. We choose *d*^*l*^ = 0.5 (*day*^−1^).

### Fitting procedure

The goal of fitting is to find *a*^*l*^and *p*^*l*^of each leukemic clone so that given the clonal percentages at diagnosis, the error of the clonal percentages at relapse is less than 1% compared to real data, subject to the constraint that BM leukemic population constitutes less than 5% of the total population in BM at the end of induction treatment (so that complete remission is achieved).

For each random parameter set, given the clonal percentages at diagnosis and the patient’s data, we calculate the clonal percentages at relapse. The “error” of the parameter set is defined as the largest element-wise difference between these and the patient’s sequencing-based clonal percentages at relapse.

The fitting scheme is defined in the following steps:

Step 1: For each clone, the initial guesses for its proliferation and renewal rates are sampled from uniform distributions:

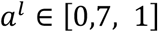

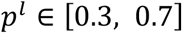
Step 2: Starting from the initial guess, use the FORTRAN procedure NELMIN (29, 30), which seeks the minimum value of a function using the Nelder-Mead algorithm, a simplex-type method, to arrive at a final guess for the parameter set that minimizes the error.
Step 3: Re-do step 2 if necessary with the newly found parameter set if the error can be reduced further.
Step 4: Check the final error. If the error is less than 1, then record the parameter set and return to step 1, until 100 parameter sets are found.

We observed that after NELMIN converges to a minimum, if we restart the procedure with the result of the old run as the initial guess for the new run, NELMIN sometimes converges to an even smaller minimum. Step 3 in the scheme above therefore makes sure that the optimization algorithm converges to a local minimum, given the random initial guess.

Note that for the clones that are present at relapse but that were not detected at diagnosis, we assume they already existed as small populations at diagnosis, instead of assuming they are new mutants. There exists evidence that resistant clones exist before treatment, instead of being driven by de novo mutations (24, 25). These clones are therefore assumed to have no mature cells in their populations and their mitotic populations occupy 3% each in bone marrow at diagnosis. This percentage was chosen to represent very small clones at diagnosis; however, the percentages can be reduced to 0.1% without changing the clonal dynamics in expected-value simulations.

### Stochastic simulation algorithm

Parameters of each leukemic clone have been determined by fitting the expected-value model to patients’ data. 1000 Monte Carlo trajectories are produced for each patient and parameter set.

For a large population of cells, Gillespie’s Stochastic Simulation Algorithm (SSA) (12) is too slow. For our case here, where the total number of cells can be in the range of 10^12^ − 10^13^, even the *τ*-leaping algorithm does not perform fast enough. We used the following decision tree:

- 10^0^ – 10^2^ cells or less each: apply SSA (12).
- 10^2^– 10^6^ cells: apply *τ*-leaping algorithm (12, 13).
- 10^6^ cells or more: apply deterministic algorithm (MATLAB’s ode23, an ODE solver, is used here).

The resulting algorithm faithfully models the stochastic effects while performing much more effectively.

### Minimal residual disease (MRD)

In addition to simulations of therapy effects, we need to derive expressions that allow us to compare our results to the independent measurements of MRD in (15). Technically, this consists of deriving an expression for the net growth rate of each leukemic clone given its proliferation and differentiation rates. The net growth rates are needed to compare the model predictions with the MRD data in (15):

1. The ODE governing the population of leukemic mitotic cells in bone marrow is

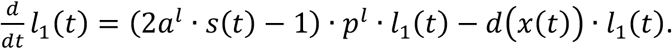
2. Negative feedbacks *s*(*t*) and *d*(*x*(*t*)) can be estimated by considering only the normal cells in blood and bone marrow (which dominate between remission and relapse). The corresponding estimates are denoted 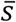 and 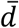, respectively.
3. The ODE now has solution

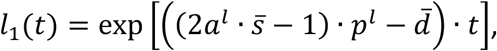

from which we derive

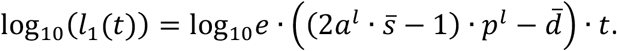
4. Adjusting the units (the parameters on the right hand side are in day^−1^ but the growth rates in (15) are in month^−1^), we conclude that

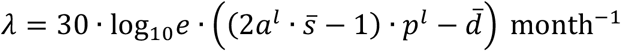

with 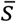 and 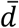 computed immediately after treatment in the expected-value solution (assuming the normal clone dominates until relapse).

## Acknowledgements

MK and SJC were supported by NIH R01HL128173, KD and SJC were supported by Alex’s Lemonade Stand Foundation Innovation Award. MK was supported by the grant 2012/04/A/ST7/00353 and RJ by the grant 2016/23/D/ST7/03665 from the National Science Center (Poland).

## Supporting information

**S1 File. Supplemental information appendix.** This contains detailed patients characteristics, clonal analysis and details from fitting the expected-value model and simulating the stochastic model.

**S1 Table. Data about the patients and the input data for the expected-value and stochastic models.** This includes characteristics, disease, nestedness of subclones and chemotherapy treatment.

**S2 Table. Parameter sets from fitting the expected-value model for all patients, 100 parameter sets per patient.**

**S1 Fig. Results of the expected-value model for Patient 1 (ID: 400220).** ***(A-D):** Results of fitting the expected-value model using a single parameter set*. **(A)** Evolution of mitotic populations in the BM of all clones in logarithmic scale. **(B)** Evolution of the mature populations in blood. Green bars indicate chemotherapy treatments. **(C)** Evolution of BM cellularity. Parameter sets are chosen so that the BM cellularity is reduced to approximately 15 – 20% of the normal value, as experimentally observed (read (31)). **(D)** Evolution of clonal percentages. The bar-plot on the left consists of clonal percentages at diagnosis, and the one on the right consists of clonal percentages at relapse (from data). The parameter sets are chosen to fit the clonality data in these two bar-plots, as shown in the middle plot. The color code for different clones in (B) and (D) is the same as in (A), described in its legend.

**S2 Fig. Results of the expected-value model for Patient 2 (ID: 426980).**

**S3 Fig. Results of the expected-value model for Patient 4 (ID: 573988).**

**S4 Fig. Results of the expected-value model for Patient 5 (ID: 758168).**

**S5 Fig. Results of the expected-value model for Patient 6 (ID: 804168).**

**S6 Fig. Results of the stochastic model for Patient 1 (ID: 400220).** Columns correspond to different parameter sets. Row 1: Evolution of clonal percentages in the expected-value model. Rows 2 and on: Outcomes of the stochastic model, in logarithmic scale with corresponding frequencies of occurrence listed.

**S7 Fig. Results of the stochastic model for Patient 3 (ID: 452198).**

**S8 Fig. Results of the stochastic model for Patient 4 (ID: 573988).**

**S9 Fig. Results of the stochastic model for Patient 6 (ID: 804168).**

**S10 Fig. Clonal evolutions for all patients.**

