## Supplemental File 1 for "Predicting Minimal Residual Disease in Acute Myeloid Leukemia through Stochastic Modeling of Clonality"

**SI appendix**

**Modeling of stochastic clonal effects in relapses following AML chemotherapy:**

**Robustness of the deterministic outcome compared to stochastic scenarios**

Khanh Dinh

Department of Mathematics, University of Alabama, Tuscaloosa, AL, USA

Roman Jaksik

Systems Engineering Group, Silesian Tech, Gliwice, Poland

Marek Kimmel

Departments of Statistics and Bioengineering, Rice University, Houston, TX, USA

Systems Engineering Group, Silesian Tech, Gliwice, Poland

**Patient characteristics**

Data about the patients, their disease and treatment from [1]:

Patient 1 (UPN ID 400220): A 34 year old Caucasian female with a prior history of cervical carcinoma in situ (treated with LEEP procedure), mitral valve prolapse, and endometriosis, presented with fatigue, allergy symptoms and chest pain. A CBC demonstrated WBC 14,500 cells/mcl, hemoglobin 11.5 g/dl, platelets 139,000 cells/mcl, with 58% circulating blasts. A bone marrow biopsy demonstrated 90% cellularity, with 71% myeloblasts (MPX +, NSE -, CD13/33/34/117/HLA-DR+), consistent with a diagnosis of AML M4. Cytogenetics showed a 46 XX karyotype. Molecular diagnostic studies demonstrated a FLT3 internal tandem duplication. Initial therapy consisted of 7 +3 induction regimen with infusional cytarabine and idarubicin. A mid-cycle bone marrow biopsy revealed ablation, and a subsequent biopsy documented first complete remission. Post-remission therapy consisted of 3 cycles of high dose cytarabine. Relapse was documented 8 months after initial diagnosis. The patient underwent salvage chemotherapy with mitoxantrone, etoposide, and high dose cytarabine (MEC) with concurrent plerixafor, but did not achieve remission. She was subsequently treated with fludarabine, high dose cytarabine, idarubicin, and gemtuzumab with concurrent filgrastim, and subsequently underwent matched unrelated donor stem cell transplantation with active disease. She expired at day +7 post transplant, 10 months from initial diagnosis.

Patient 2 (UPN ID 426980): A 69 year old Caucasian male, with a prior history of hypothyroidism, hyperlipidemia, and pneumonia, presented with low grade fever and cough. A CBC demonstrated WBC 6000 cells/mcl, hemoglobin 7.4 g/dl, platelets 102,000 cells/mcl, with 29% circulating blasts. A bone marrow biopsy demonstrated 50% cellularity, with 64% myeloblasts (MPX +, NSE -, CD13/33/117+), consistent with a diagnosis of AML M2. Cytogenetics showed a 46 XY karyotype. Initial therapy consisted of 7 + 3 induction with infusional cytarabine, daunorubicin, and concurrent oblimersen (Genasense, a BCL2 antisense molecule). A mid-cycle bone marrow biopsy revealed ablation, and a subsequent biopsy documented first complete remission. Post-remission therapy consisted of 2 cycles of high dose cytarabine with concurrent oblimersin. Relapse was documented 26 months from diagnosis. Salvage chemotherapy regimens were administered sequentially with no or minimal response including decitabine, high dose cytarabine, mitoxantrone and etoposide, azacytidine, and palliative hydroxyurea. He expired from progressive disease 48 months following his initial diagnosis.

Patient 3 (UPN ID 452198): A 55 year old Caucasian male, previously healthy, presented with fatigue, subjective fevers, and sinus congestion. A CBC demonstrated: WBC 72,600 cells/mcl, hemoglobin 8.2 g/dl, platelets 17,000 cells/mcl, with 8% circulating blasts and 50% monocytes. A bone marrow biopsy was inevaluable for cellularity. The aspirate demonstrated 97 % monoblasts (MPX -, NSE +, CD13/33/64+), consistent with a diagnosis of AML M5. Cytogenetics showed a 46 XY karyotype. Initial therapy consisted of hydroxyurea for two days, followed by 7 + 3 induction with infusional cytarabine and idarubicin. A mid-cycle bone marrow biopsy revealed ablation, and a subsequent biopsy documented first complete remission. Post-remission therapy consisted of 4 cycles of high dose cytarabine. Relapse was documented 16 months from diagnosis, and treated with mitoxantrone, etoposide, and high dose cytarabine (MEC) with concurrent plerixafor; the patient achieved a second complete remission. He underwent matched sibling donor stem cell transplantation following busulfan/cyclophosphamide conditioning, and remains alive and in complete remission 56 months from initial diagnosis.

Patient 4 (UPN ID 573988): A 67 year old Caucasian female with a prior history of hyperthroidism (treated with radioactive iodine), hypertension, pancreatitis, and breast intraductal carcinoma in situ (treated with excision), presented with syncope. A CBC demonstrated WBC 15,200 cells/mcl, hemoglobin 8.2 g/dl, platelets 79 cells/mcl, with 10% circulating blasts and 30% monocytes. A bone marrow biopsy demonstrated hypercellularity, with 17% myeloblasts an (MPX +, NSE -, CD13/33/117+), and 58 % monoblasts (MPX -, NSE +, CD13/33/64+) consistent with a diagnosis of AML M4. Cerebrospinal fluid was negative for malignant cells. Cytogenetics showed a 46 XX karyotype. Initial therapy consisted of 7 + 3 induction with infusional cytarabine and idarubicin. A mid-cycle bone marrow biopsy revealed ablation, and a subsequent biopsy following marrow recovery documented first complete remission. Post-remission therapy consisted of a single cycle of 5 + 2 infusional cytarabine and idarubicin. Relapse (skin and bone marrow) was documented 12 months from diagnosis and was treated with decitabine without response. She expired from progressive disease 20 months from initial diagnosis.

Patient 5 (UPN ID 758168): A previously healthy 25 year old Caucasian female presented with fatigue, nausea, vomiting, and decreased visual acuity in her left eye. A CBC demonstrated WBC 3,500 cells/mcl, hemoglobin 5.9 g/dl, platelets 24,000 cells/mcl, with circulating promyelocytes. Diffuse intravascular coagulation (DIC) was present (INR of 3.2 and fibrinogen 88 mg/dl). A bone marrow biopsy demonstrated hypercellularity, with 93% promyelocytes (MPX +, NSE -, CD13/33+), consistent with a diagnosis of AML M3. Cytogenetics showed a t(15;17) translocation. Ophthalmologic evaluation demonstrated retinal hemorrhage, detachment, and acute glaucoma, resulting in irreversible loss of vision and eventual enucleation. Initial therapy consisted of 7 + 3 induction with infusional cytarabine, idarubicin, and concurrent ATRA, complicated by headaches attributed to pseudotumor cerebri (CSF negative for malignancy). A bone marrow biopsy following marrow recovery documented first complete remission. Following induction, additional ATRA was withheld because it was presumed to be the cause of her pseudotumor cerebri. Post-remission therapy consisted of 3 cycles of "anthracycline" administered by the referring oncologist, followed by planned arsenic trioxide maintenance (prematurely aborted due to noncompliance). Relapse was documented 32 months from diagnosis, and was treated with arsenic trioxide, resulting in second complete remission. She subsequently underwent matched unrelated donor stem cell transplantation 4 months later, complicated by acute and chronic graft vs host disease and recurrent infection. She expired while in remission 50 months from initial diagnosis (14 months post transplant) from infectious complications of her transplant.

Patient 6 (UPN ID 804168): A 53 year old Caucasian male with a history of hyperlipidemia, hypertension, and coronary artery disease (treated with stent), presented with a syncopal episode. A CBC demonstrated WBC 88,100 cells/mcl, hemoglobin 8.8 g/dl, platelets 30,000 cells/mcl, with 52% circulating blasts. A bone marrow biopsy demonstrated >90% cellularity, with 86% myeloblasts (MPX+, NSE -, CD13/33/117+), consistent with a diagnosis of AML M1. Cytogenetics showed a 46 XY karyotype. Initial therapy consisted of 7 + 3 + 3 induction with infusional cytarabine, daunorubicin, and etoposide. A mid-cycle bone marrow biopsy revealed ablation, and a subsequent biopsy following marrow recovery documented first complete remission. Post-remission therapy consisted of 2 cycles of high dose cytarabine. Relapse was documented 8 months from diagnosis, and was treated with mitoxantrone, etoposide, and high dose cytarabine with concurrent plerixafor, following which residual disease was documented (5-7% blasts). He subsequently underwent matched sibling donor allogeneic stem cell transplant following conditioning with single dose total body irradiation and high dose cyclophosphamide, but relapsed 1 month post transplant. Subsequent salvage therapy included decitabine, which yielded a third remission. He subsequently developed extramedullary relapse that was treated with radiation, and died of progressive disease 30 months from initial diagnosis.

**Clonal analysis**

Clonal analysis was performed based on the variant allele frequencies (VAFs) of somatic variants (SVs) from whole-genome sequencing (WGS) of primary tumour–relapse pairs and matched skin samples from all patients [1]. The clonal nestedness is included in Supplemental table 1.

**Parameter fitting for the deterministic model**

For each patient, 100 parameter sets are found by fitting the deterministic model to the patient's information and clonal landscape at diagnosis and relapse while making sure that the fit has to satisfy the following biological constraints:

- The total leukemic population at the end of induction treatment constitutes less than 5% of the total population in the bone marrow, corresponding to the complete remission which all patients are considered having achieved.
- The BM cellularity decreases to ca. $15-20\%$ at the end of chemotherapy, which has been observed in clinic [2].

The parameter sets for all patients are given in Supplemental table 2.

**Stochastic simulation**
