## Supplementary figures and images for "Predicting Minimal Residual Disease in Acute Myeloid Leukemia through Stochastic Modeling of Clonality"

### S1_Fig.png

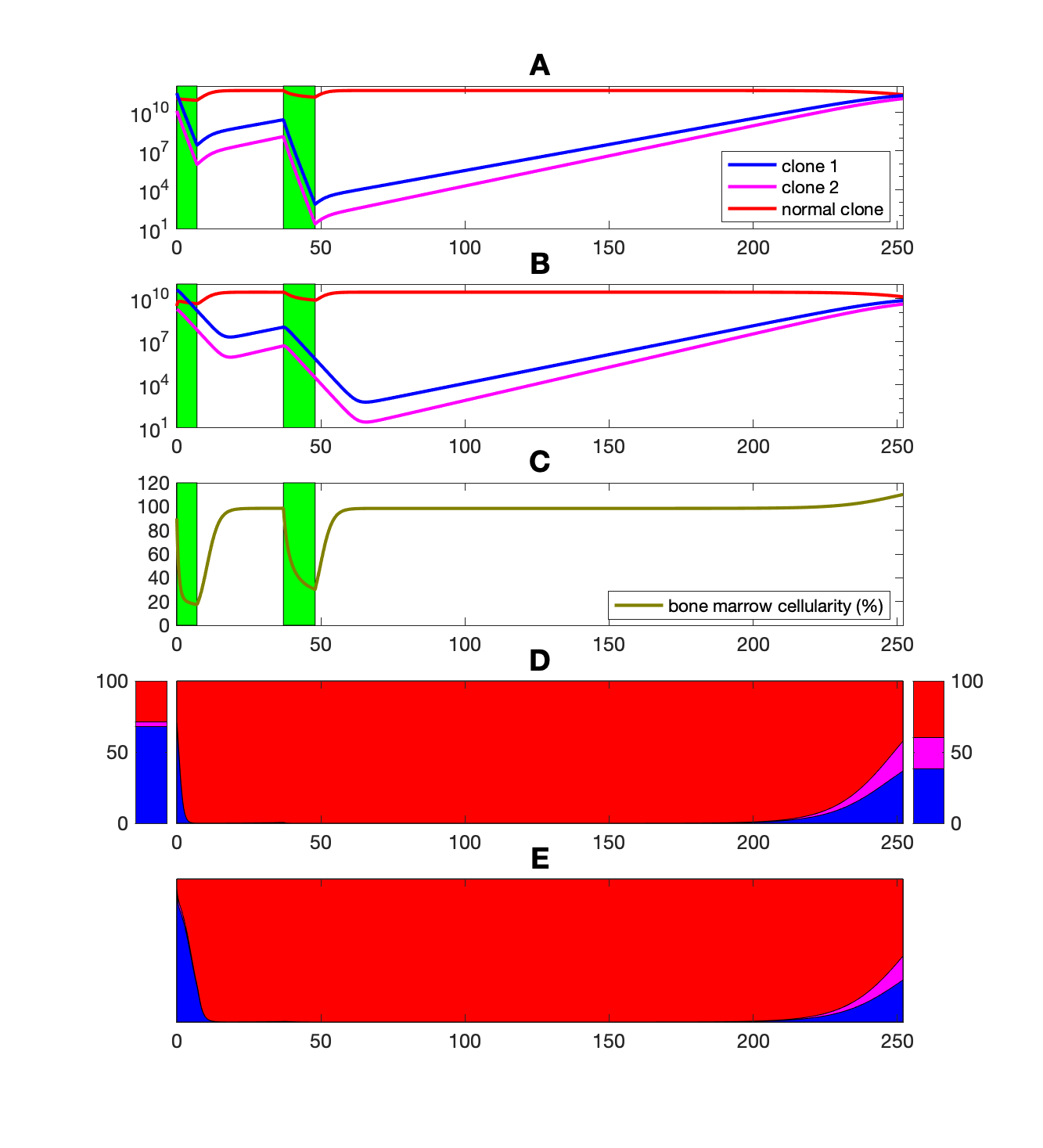

### S2_Fig.png

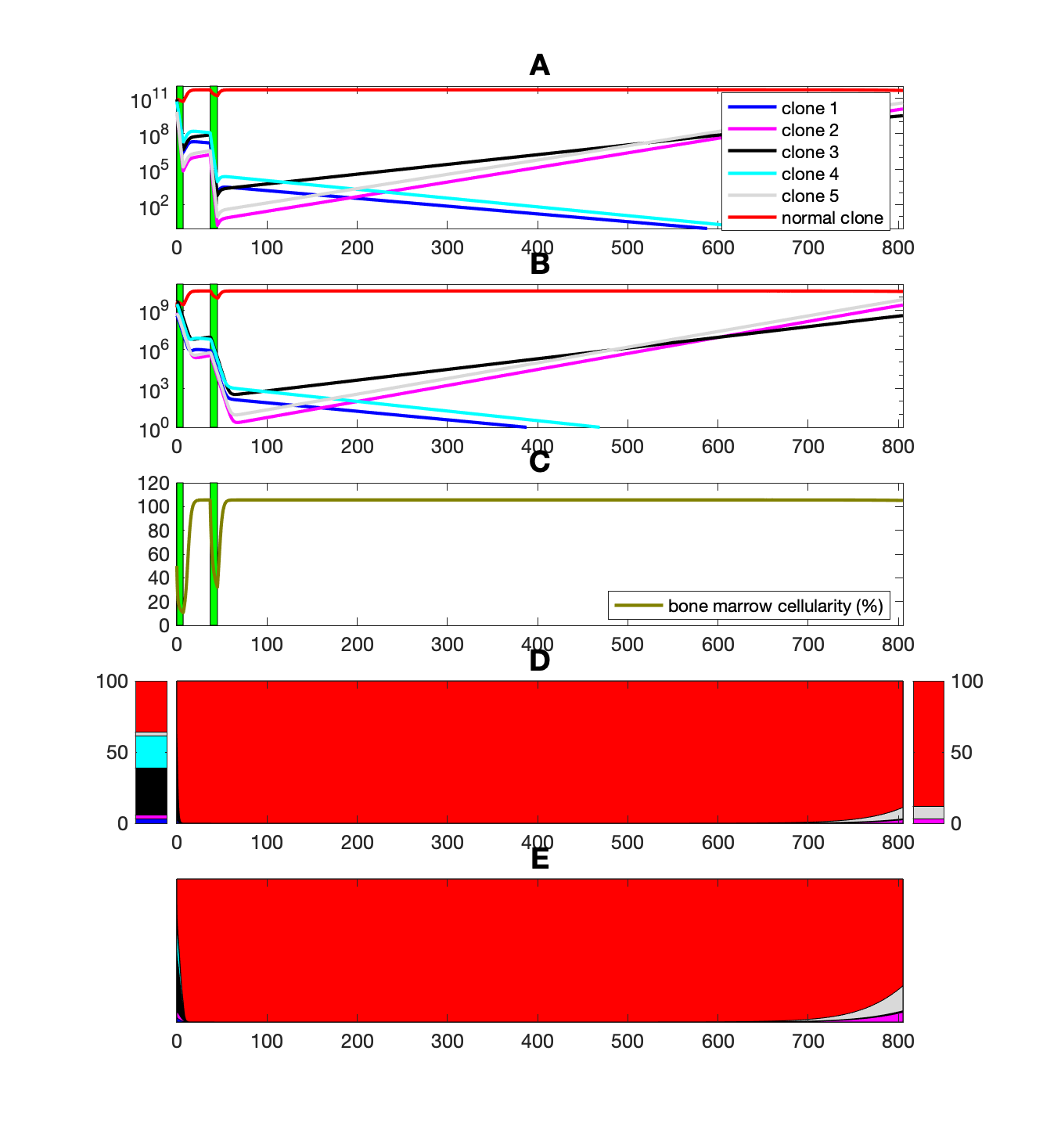

### S3_Fig.png

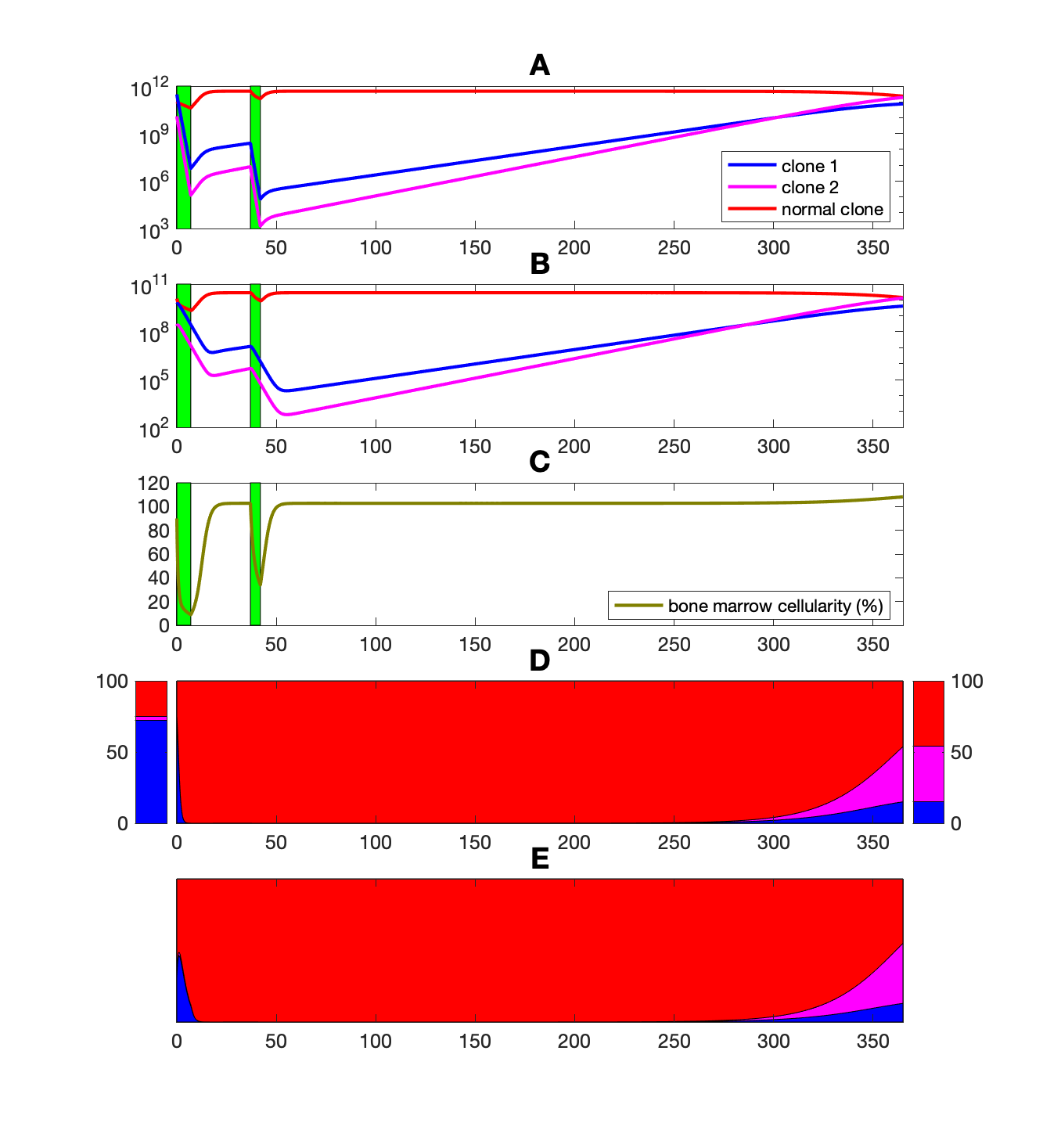

### S4_Fig.png

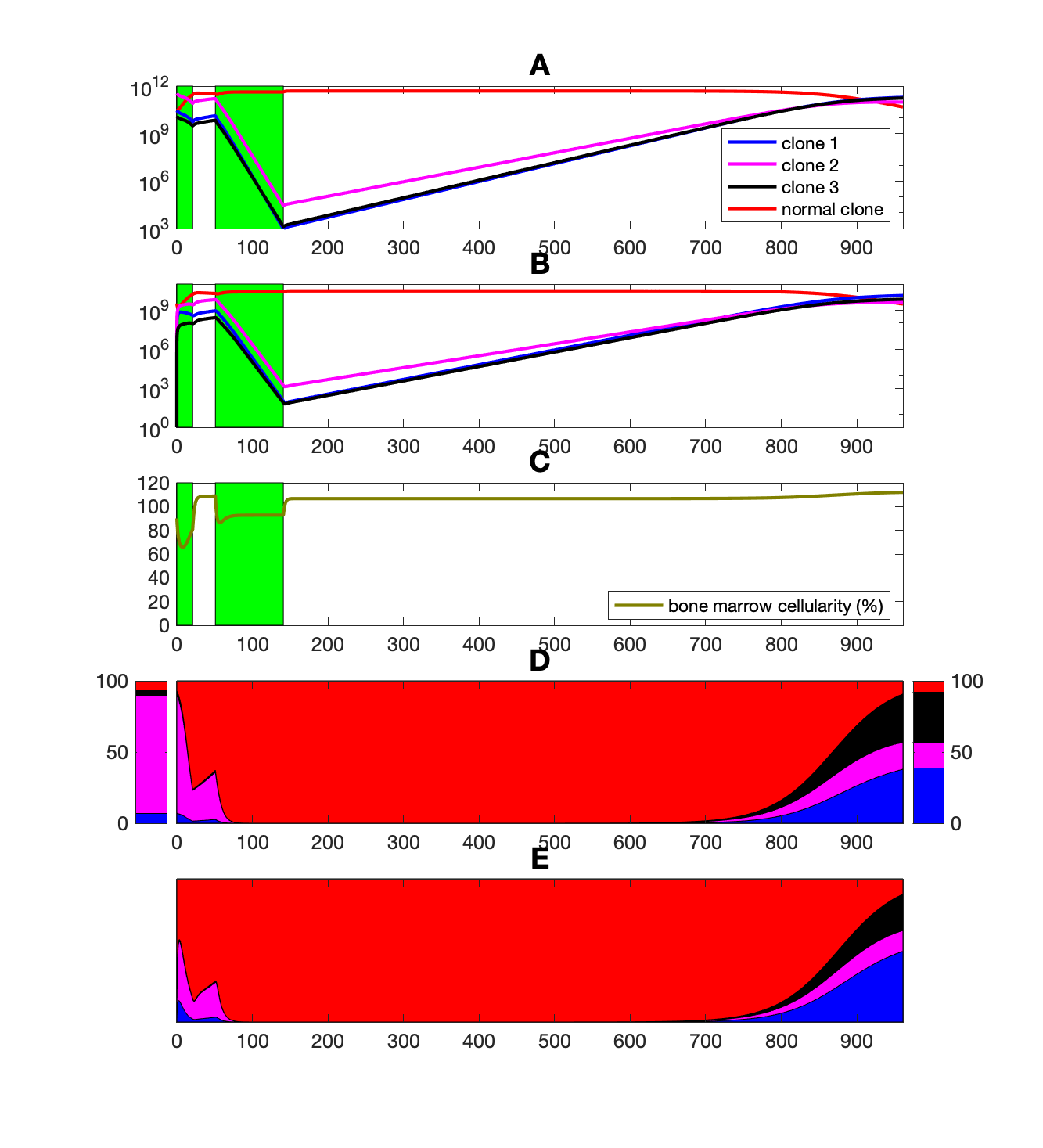

### S5_Fig.png

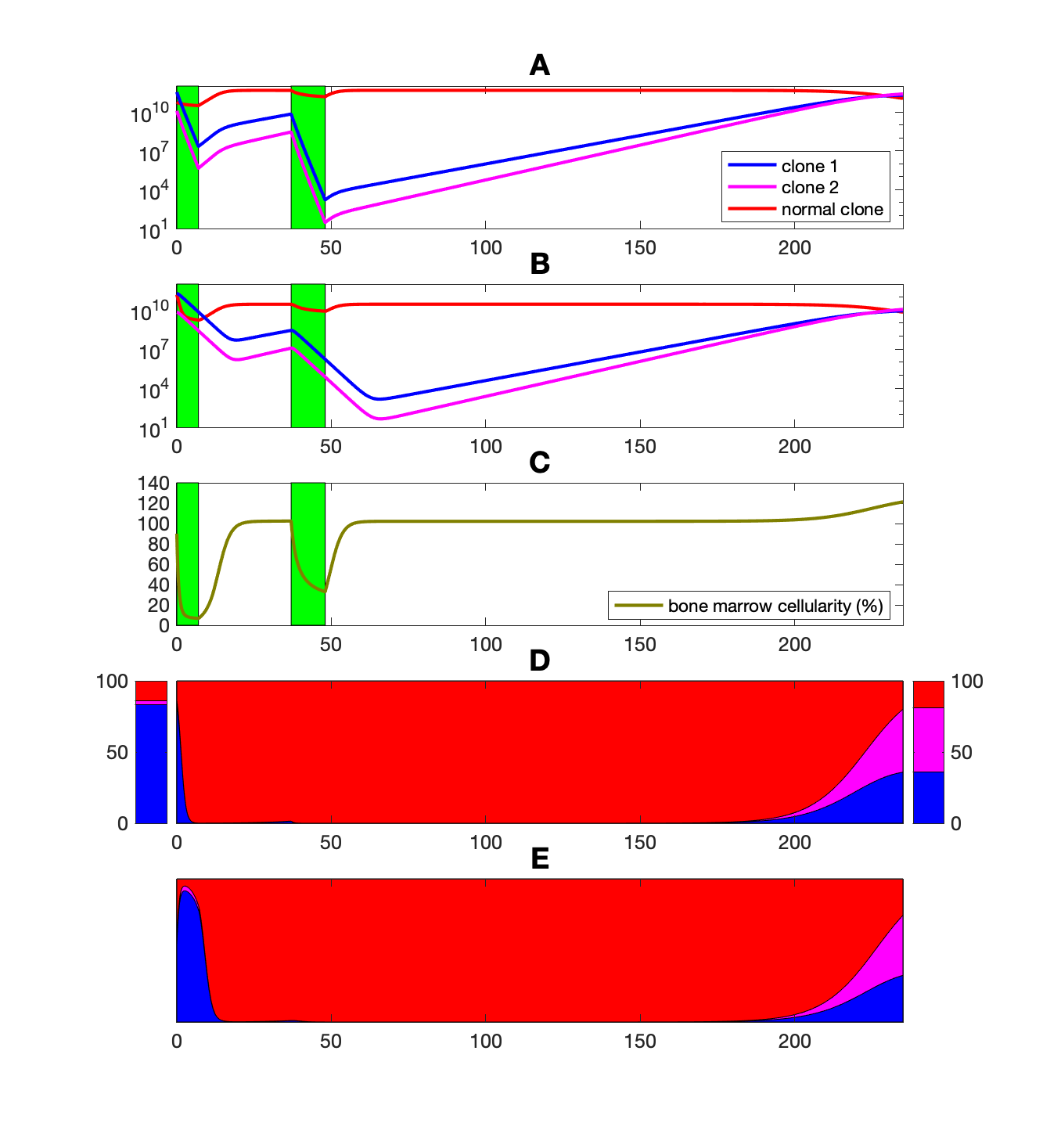

### S6_Fig.png

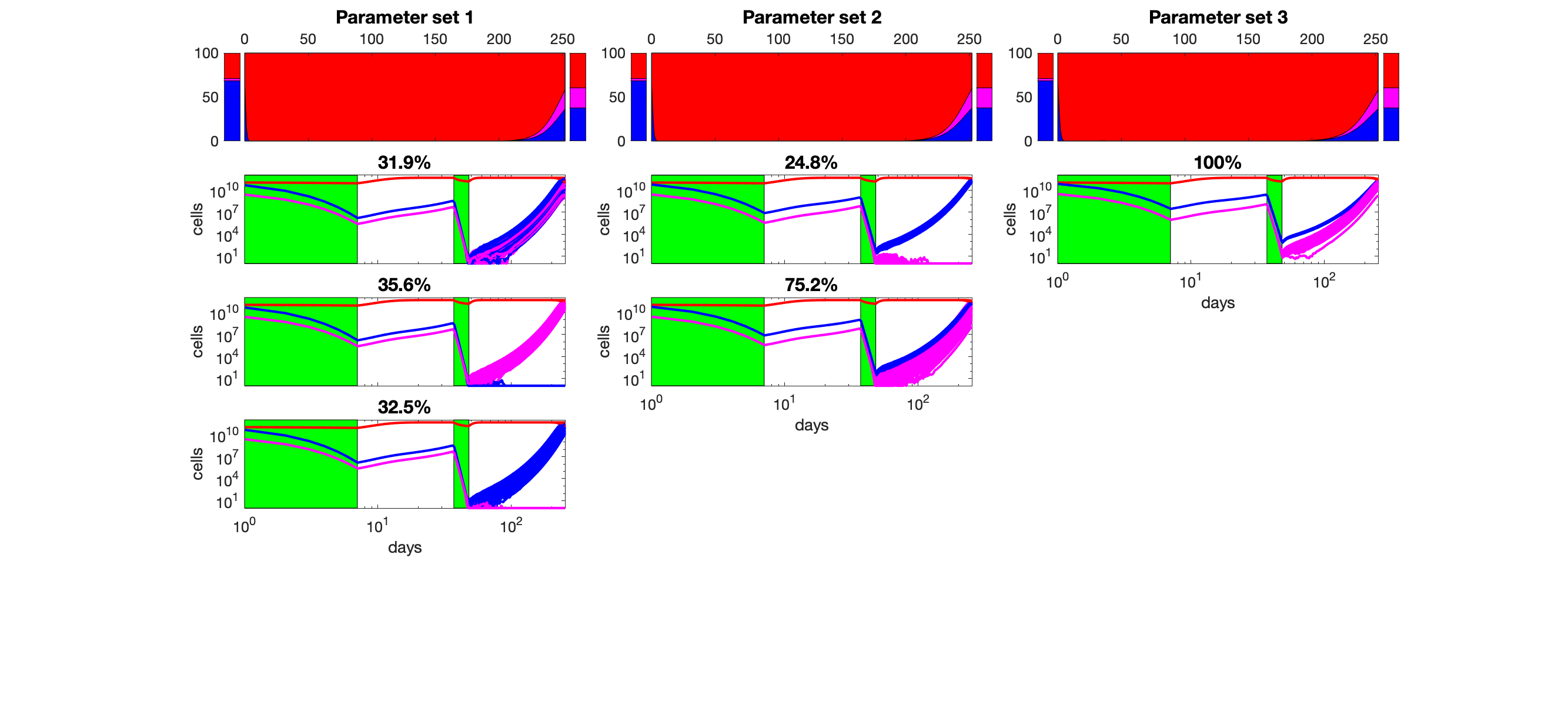

### S7_Fig.png

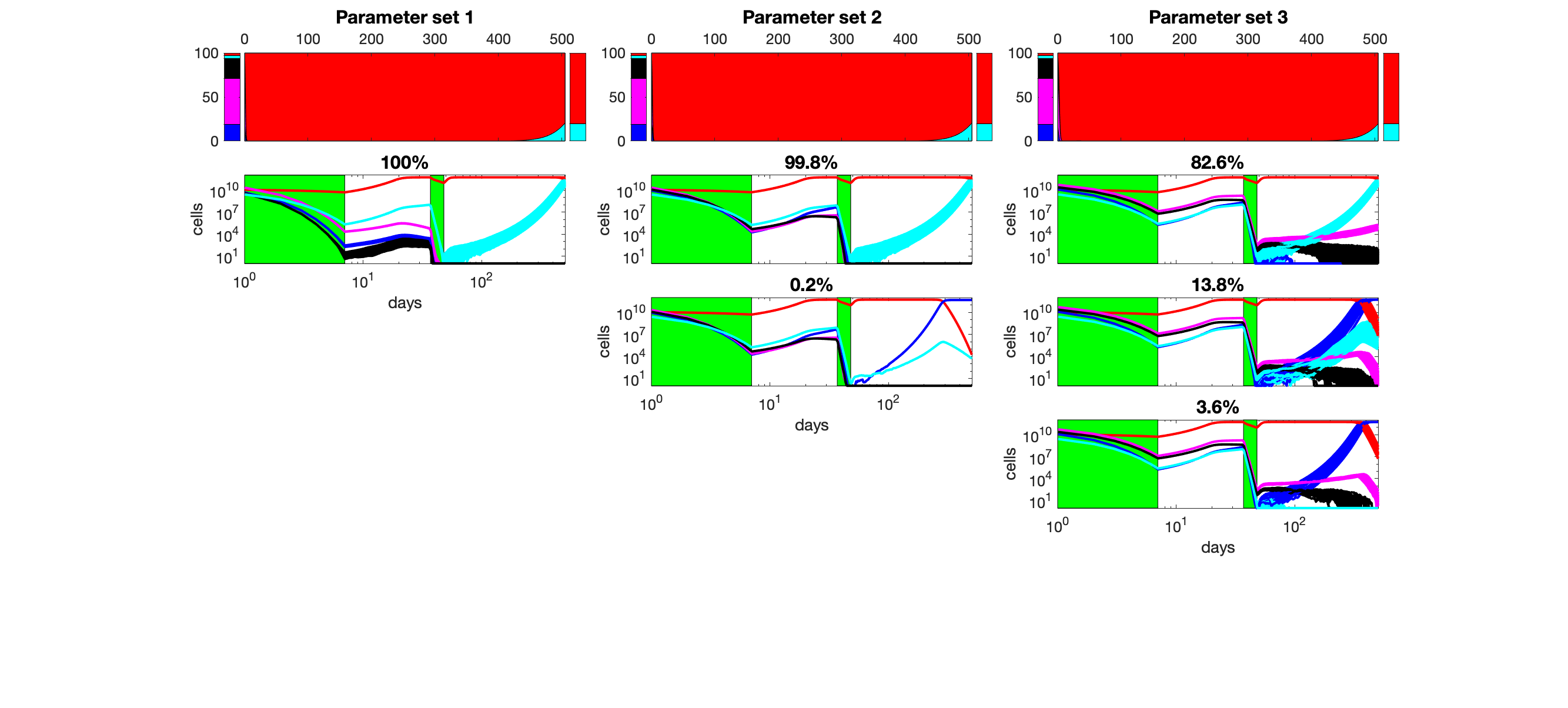

### S8_Fig.png

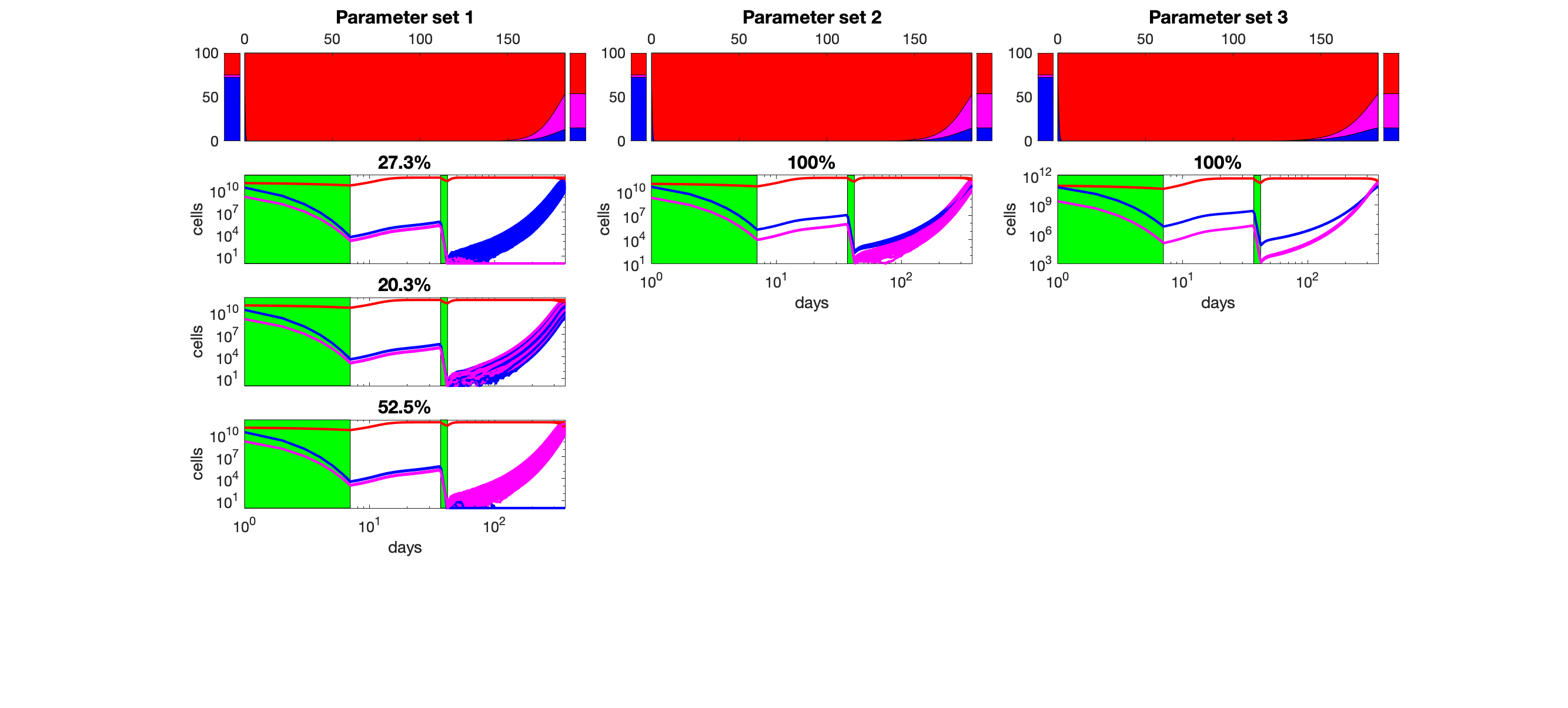

### S9_Fig.png

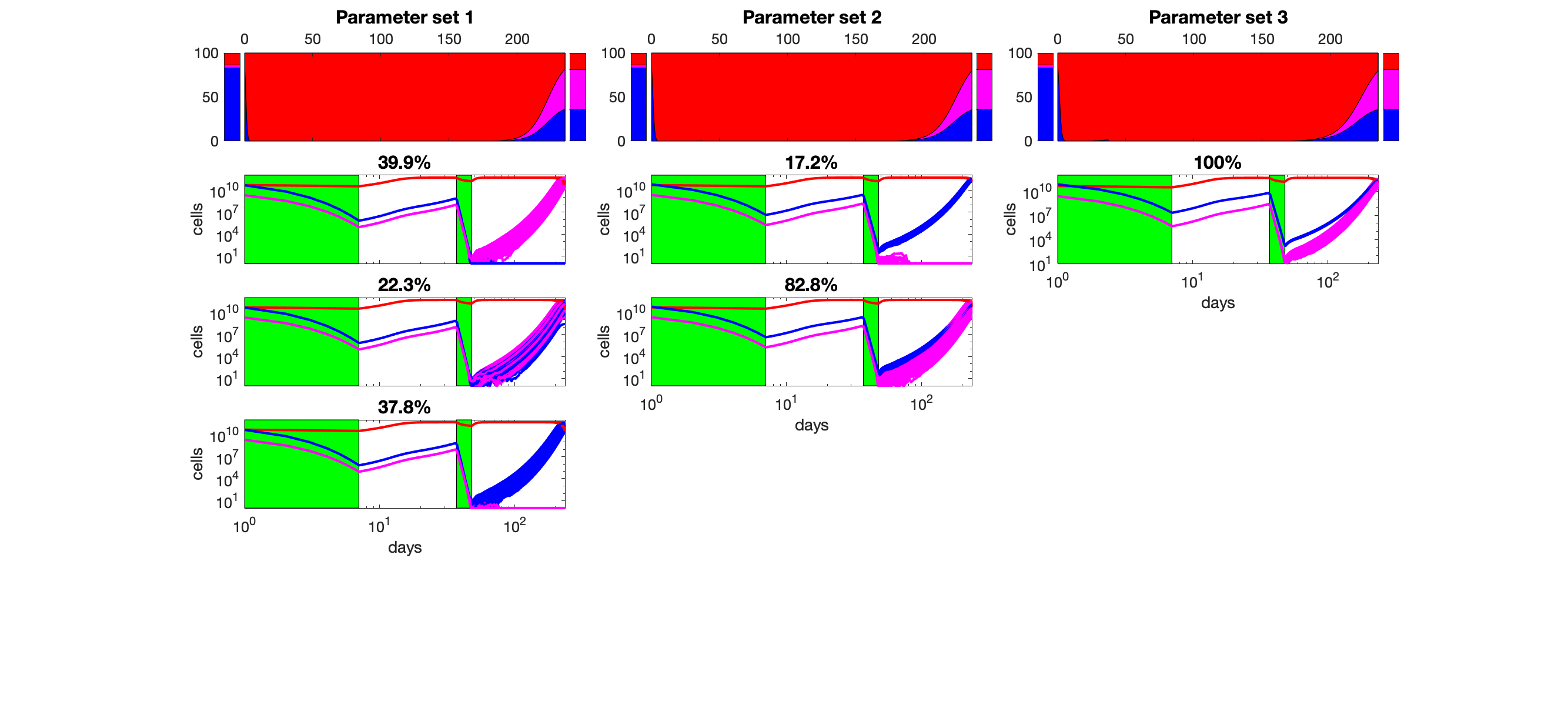

### S10_Fig.png

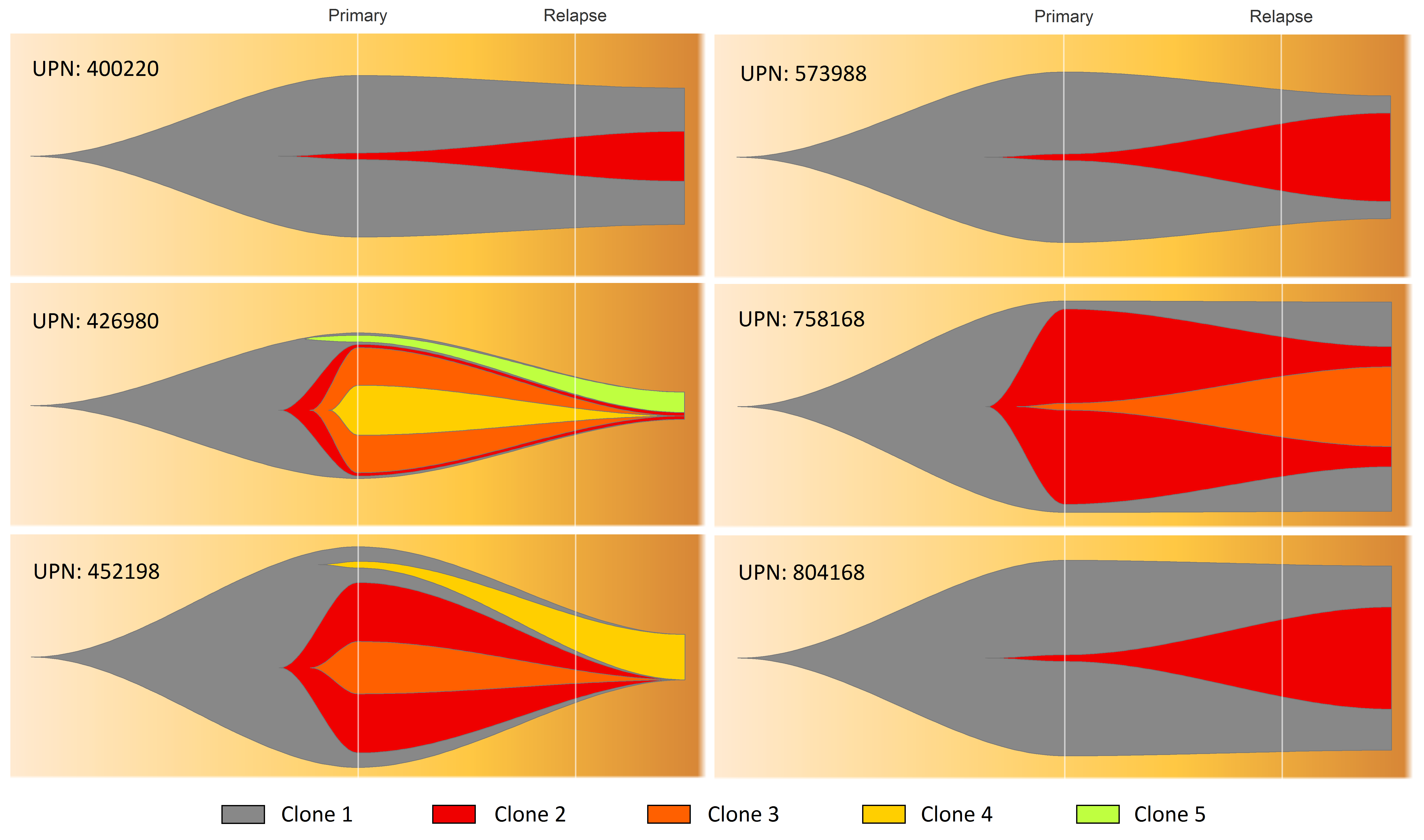

### S11_Fig.png

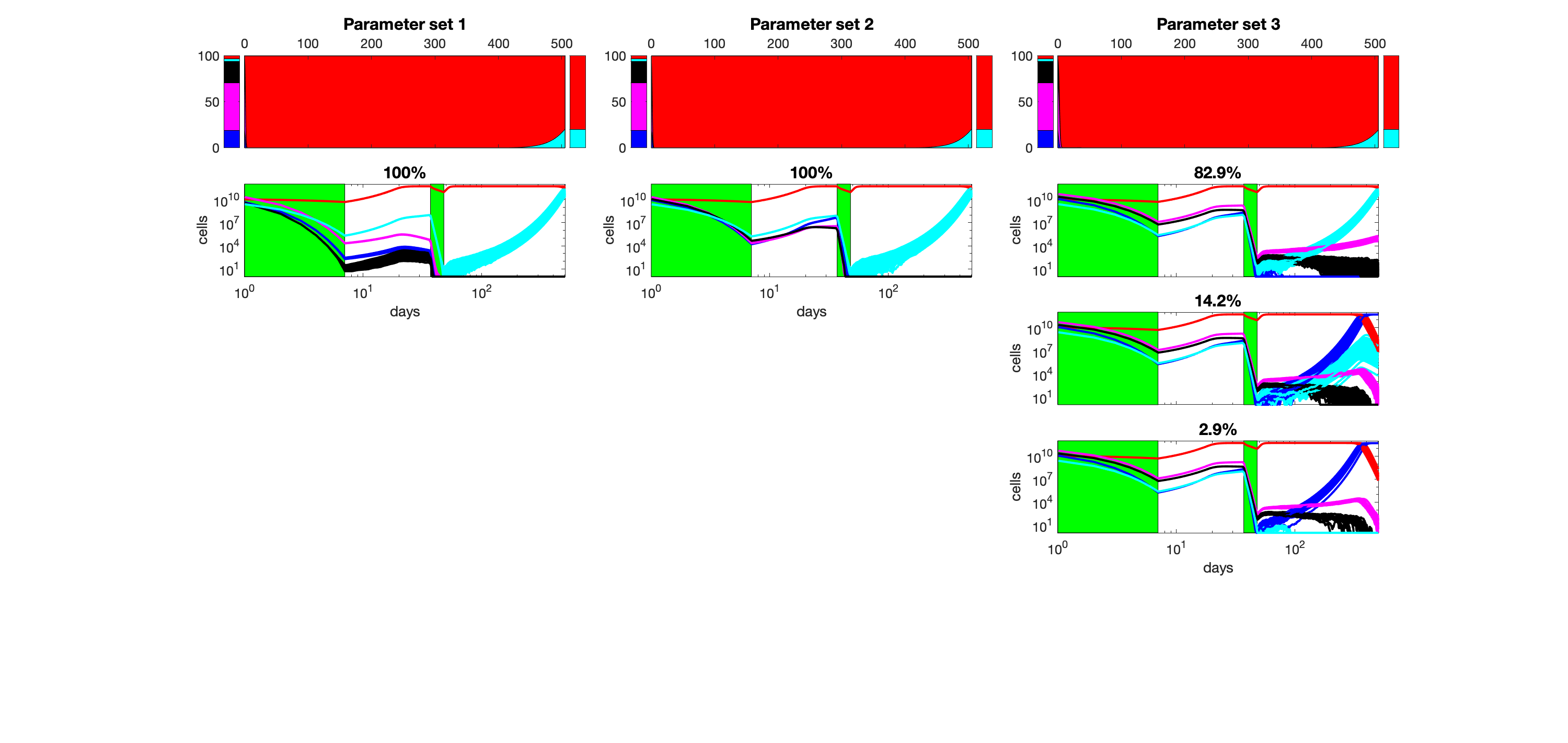

### S12_Fig.png

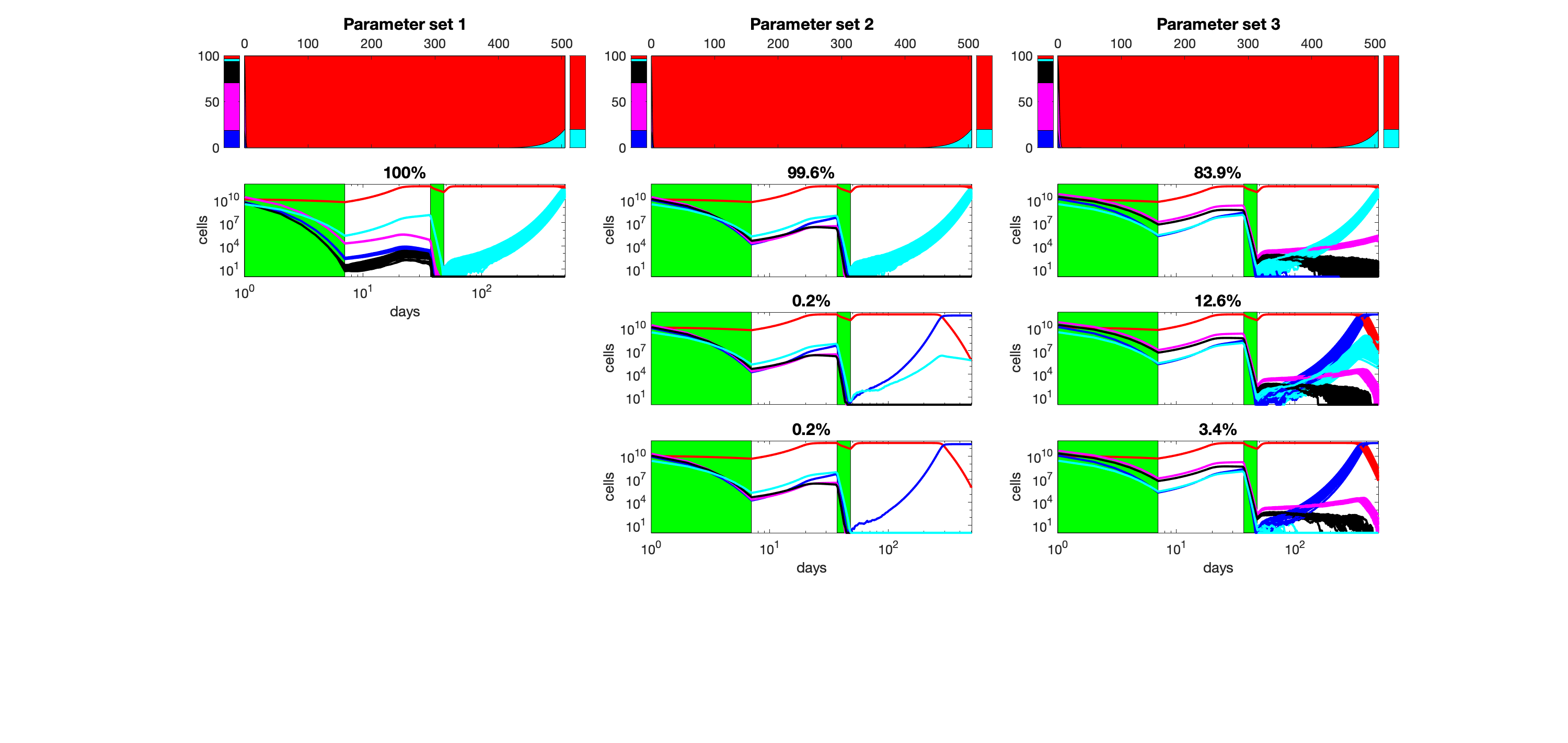

### S13_Fig.png

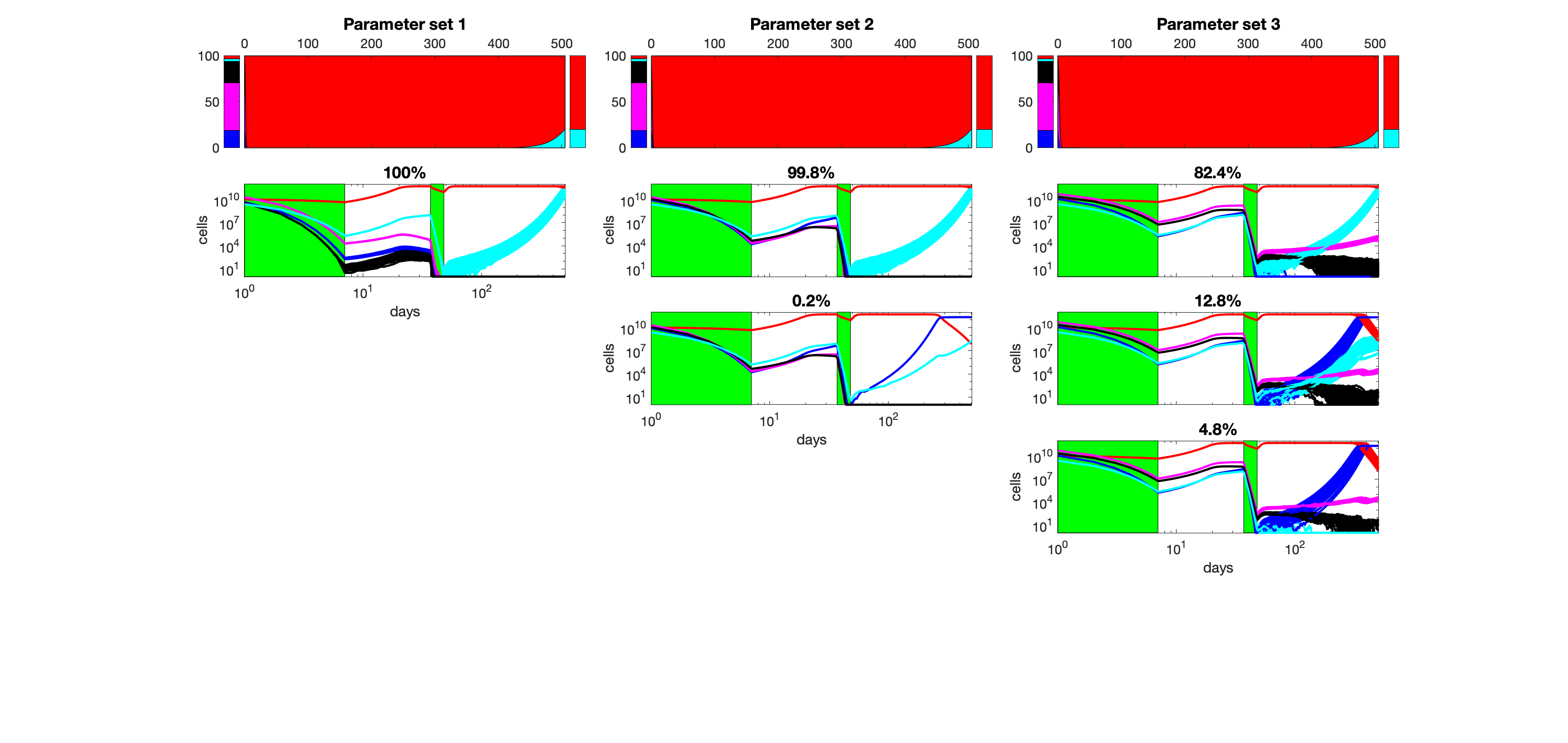

### S14_Fig.png

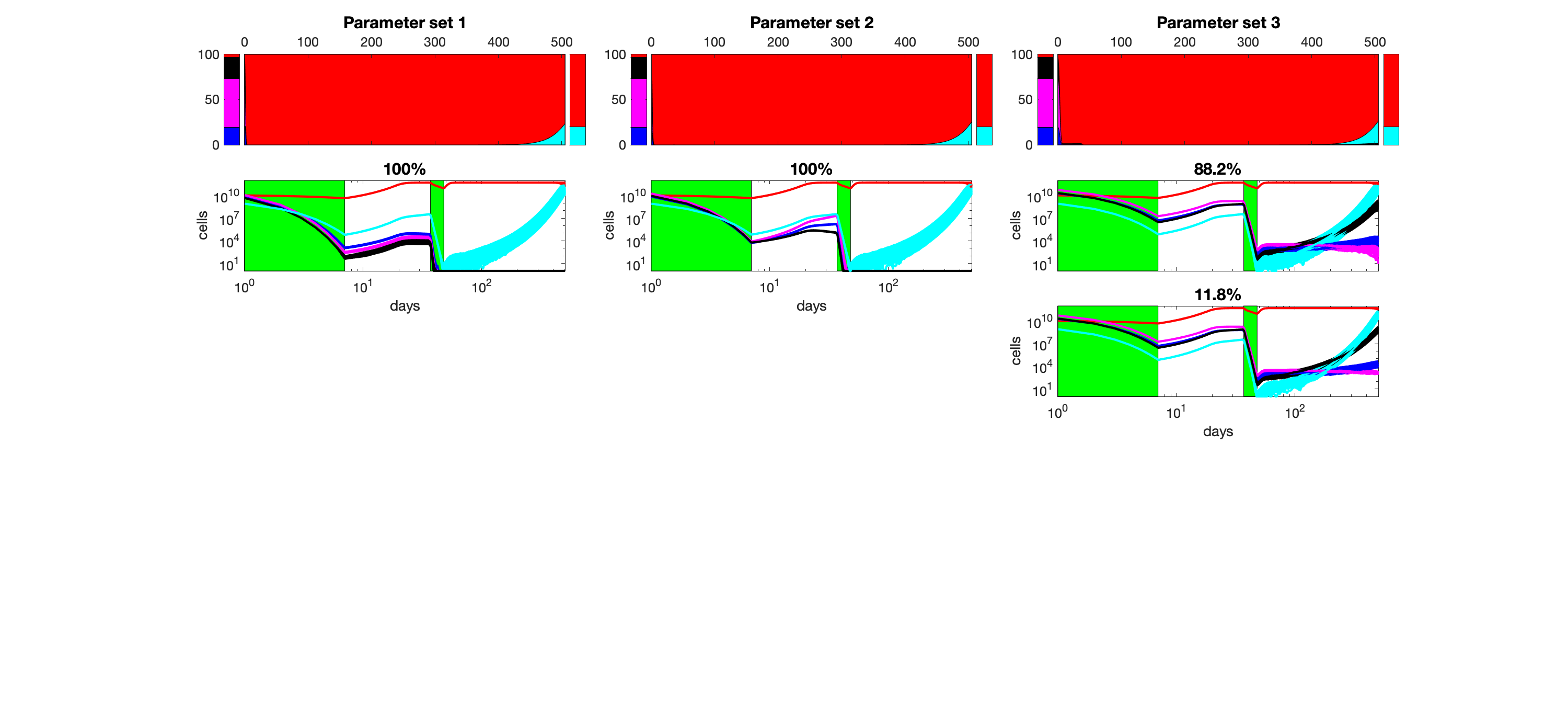

### S15_Fig.png

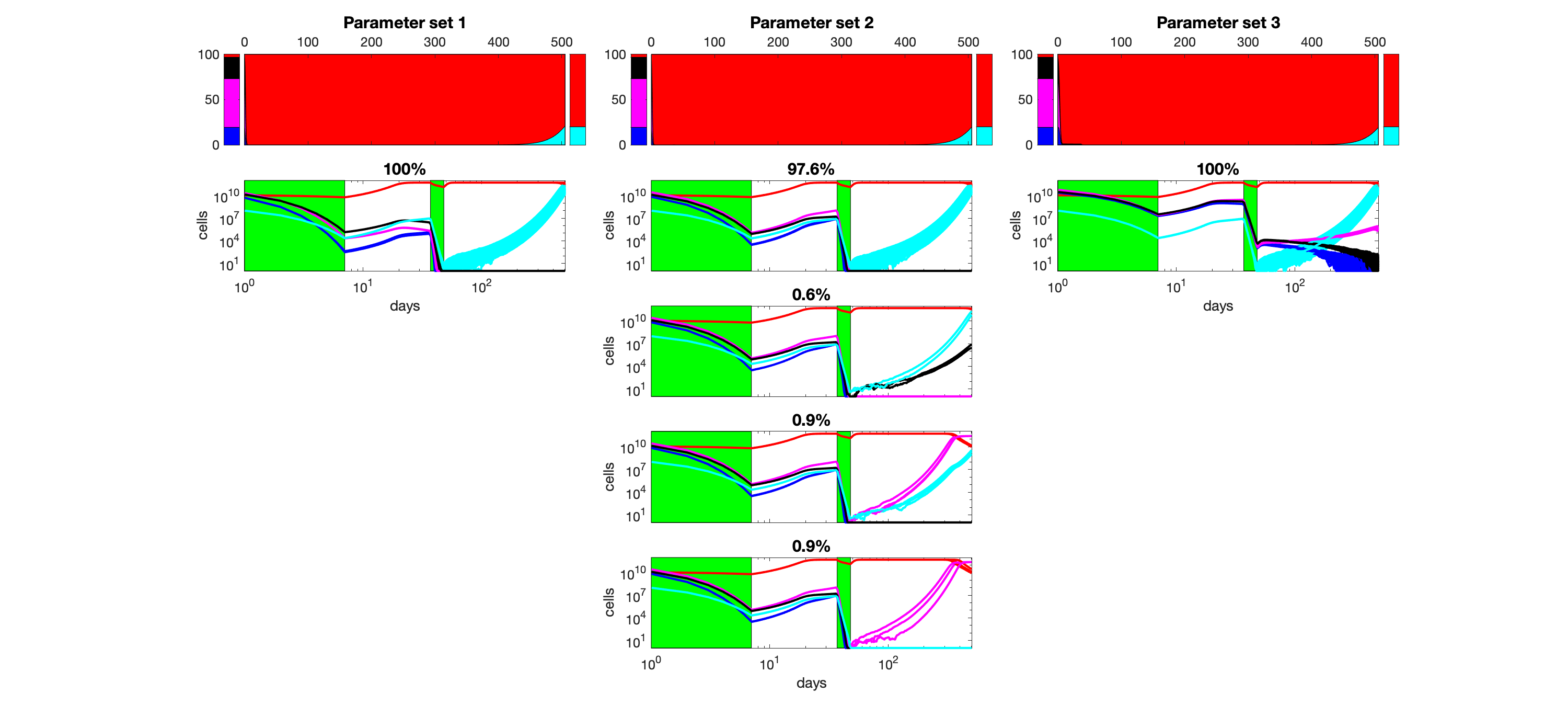

### S16_Fig.png

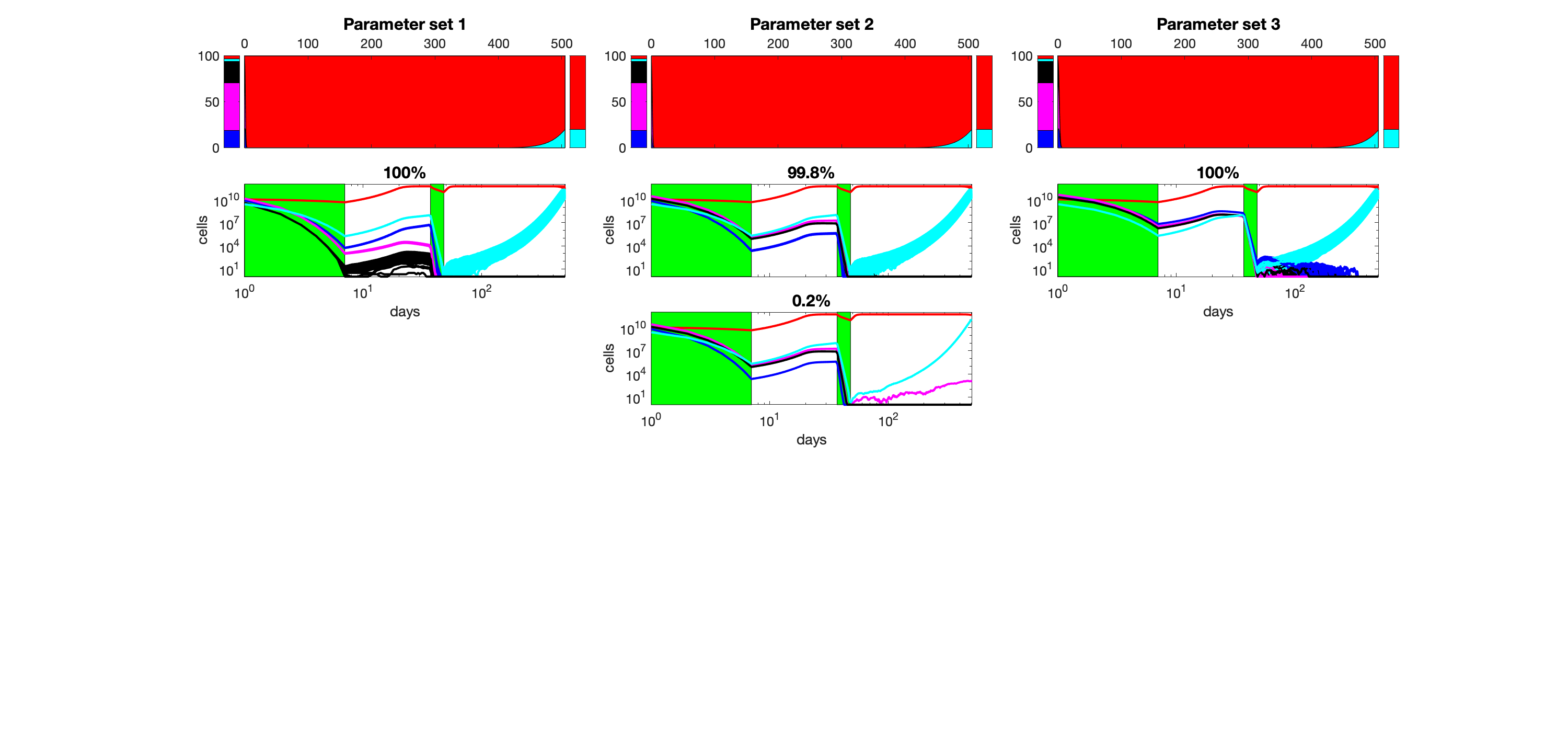

### S17_Fig.png

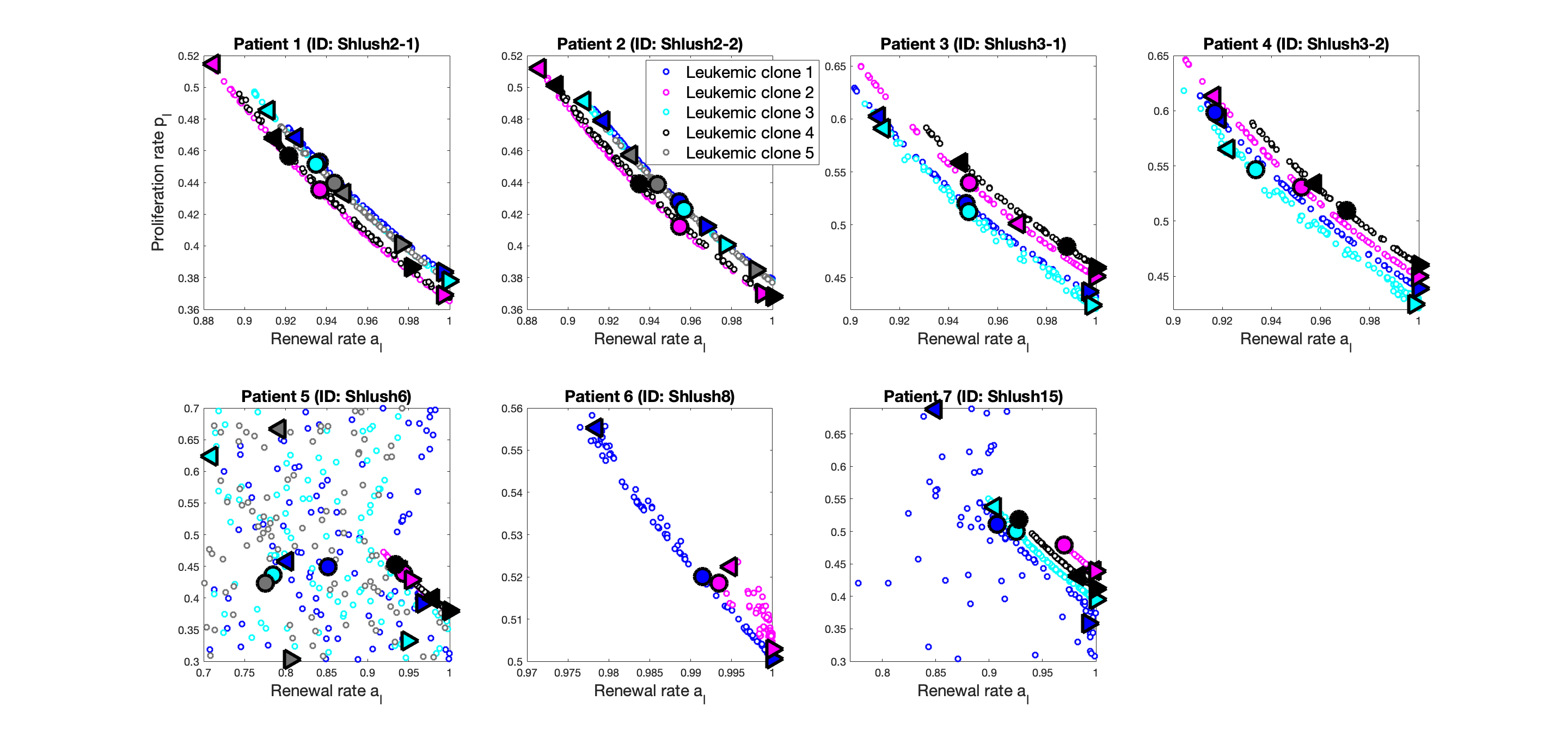

### S18_Fig.png

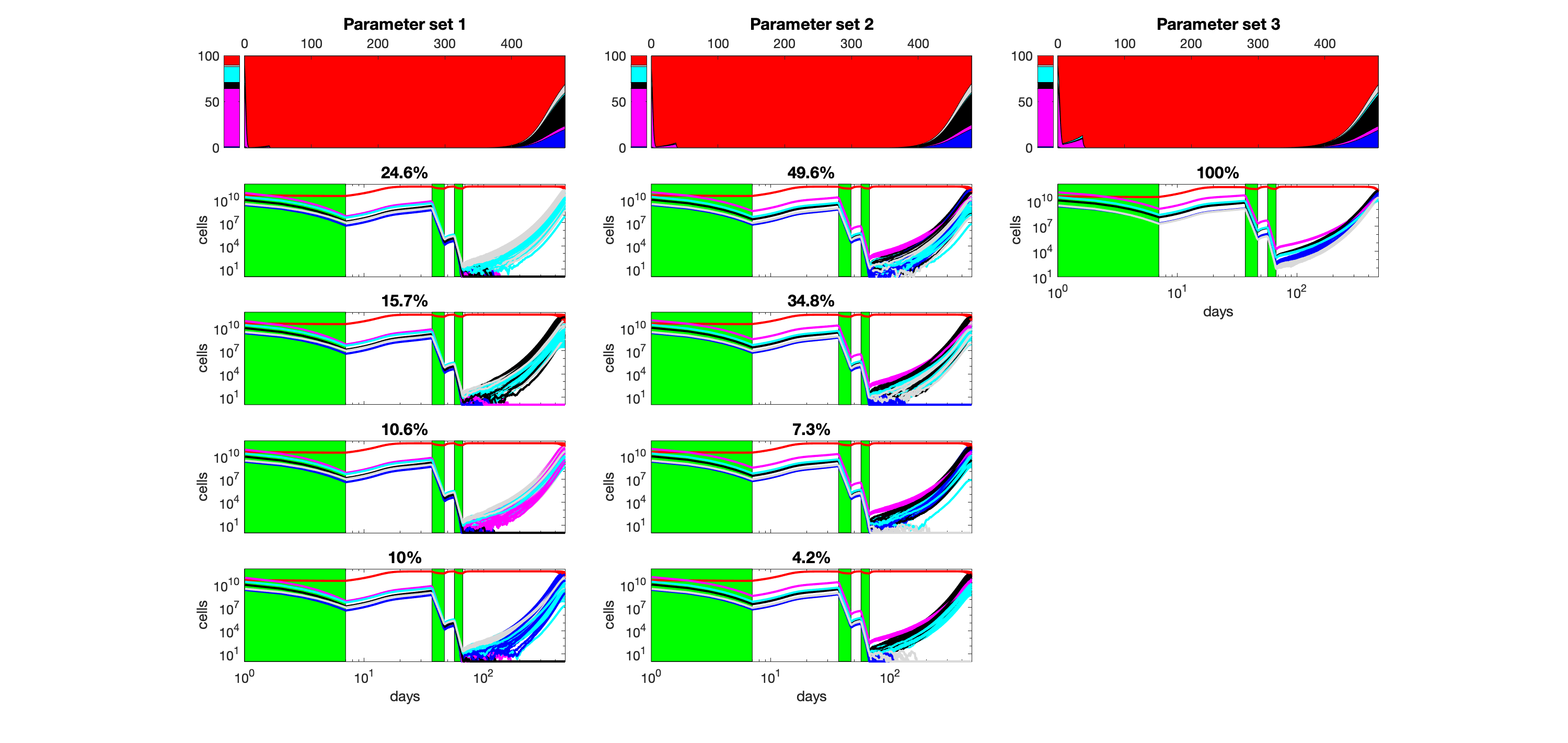

### S19_Fig.png

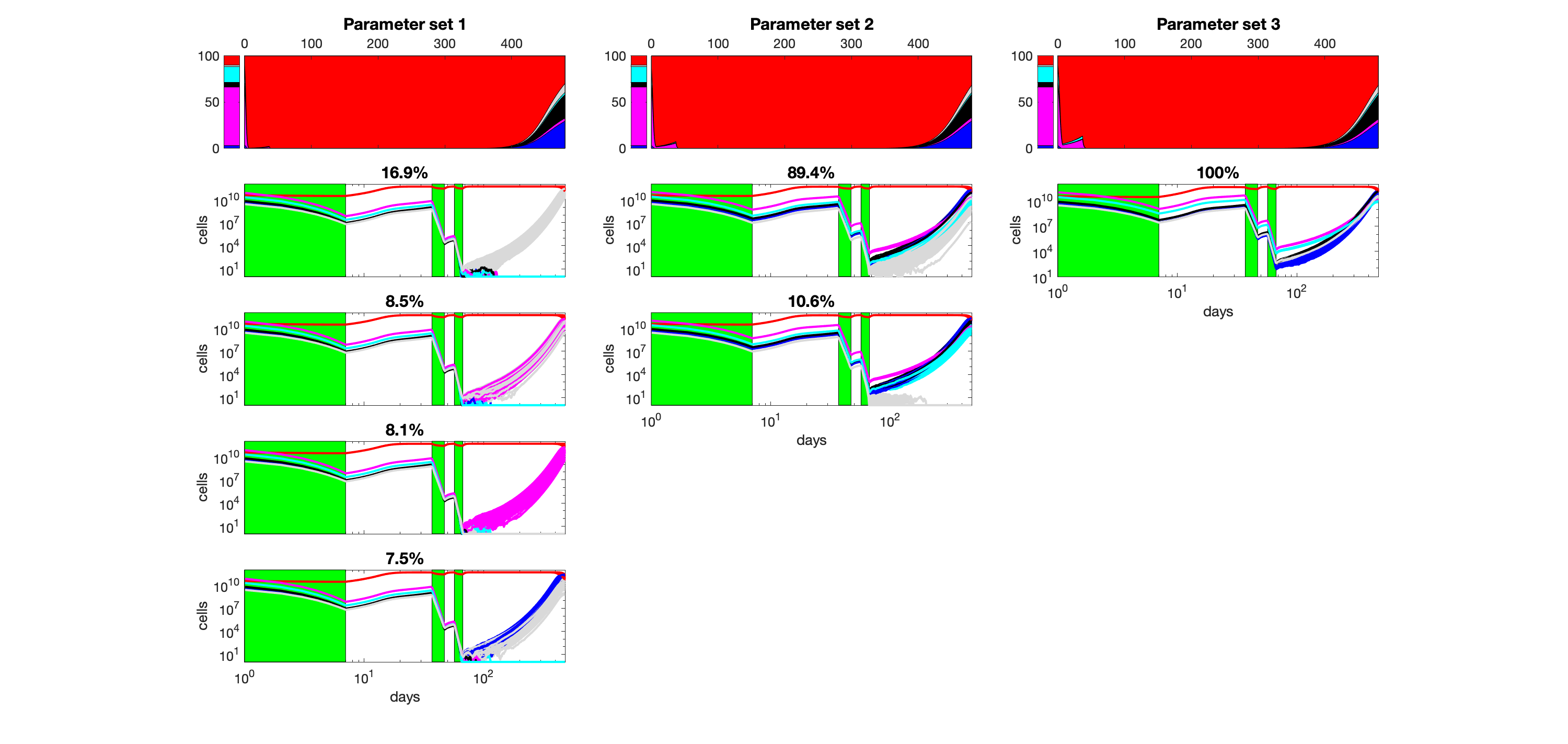

### S20_Fig.png

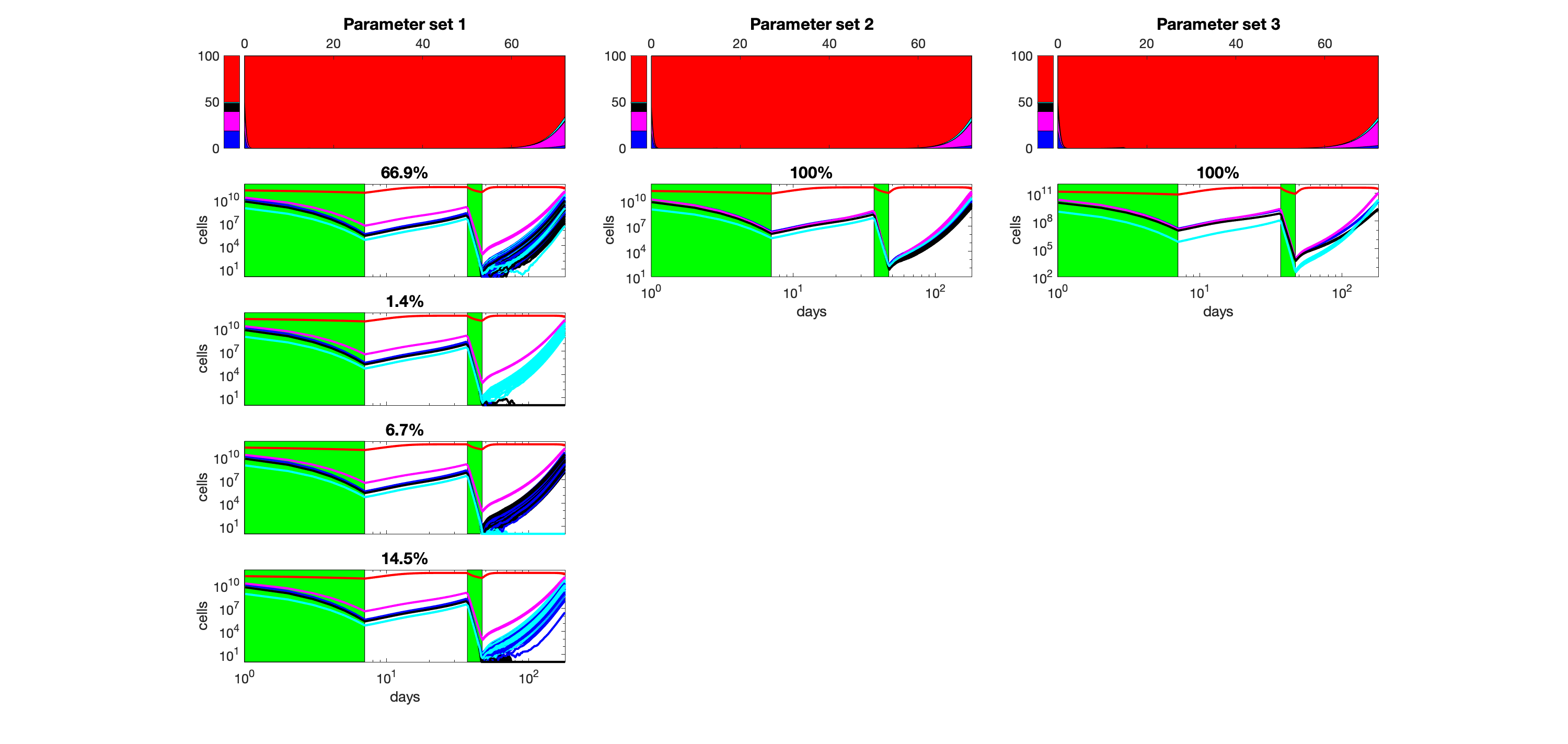

### S21_Fig.png

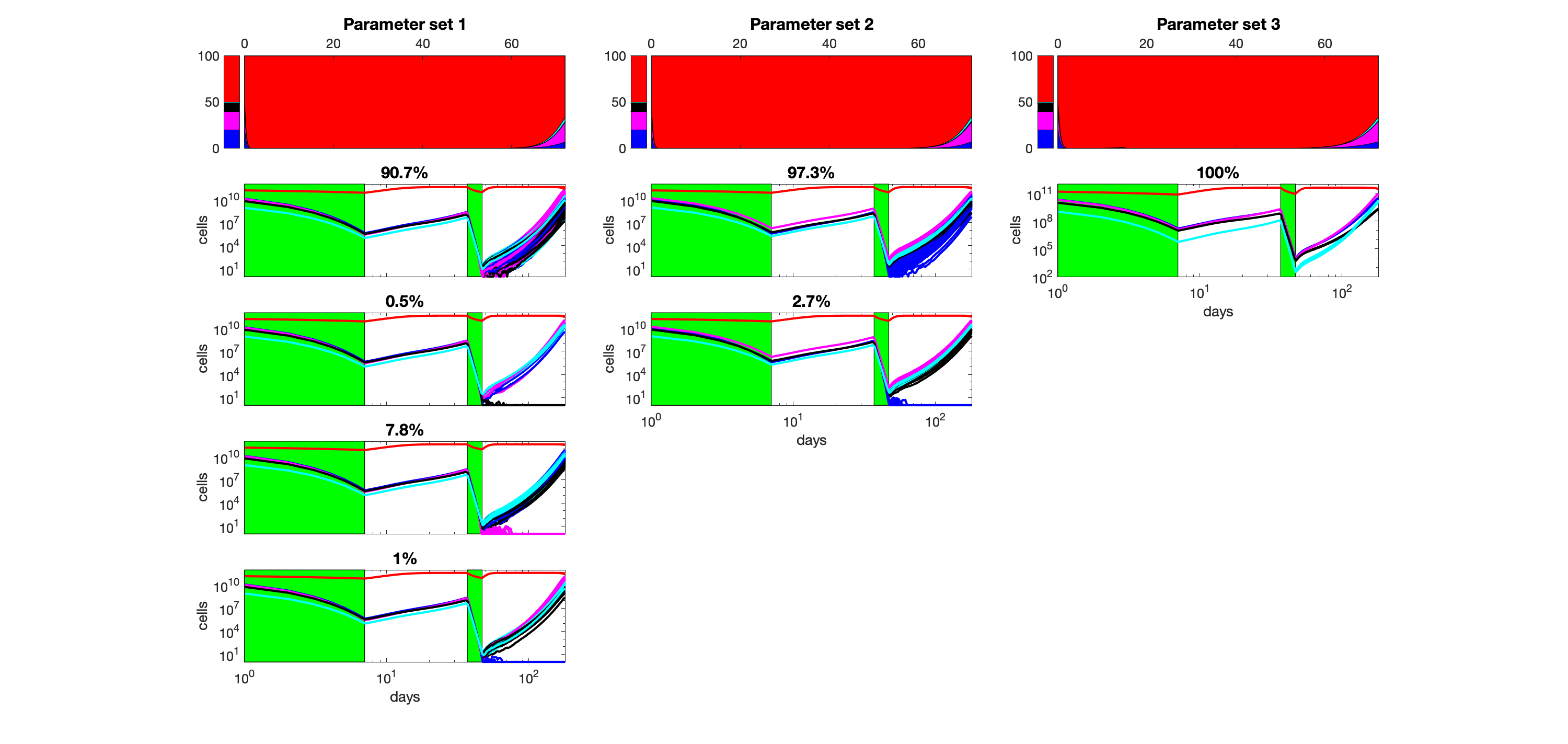

### S22_Fig.png

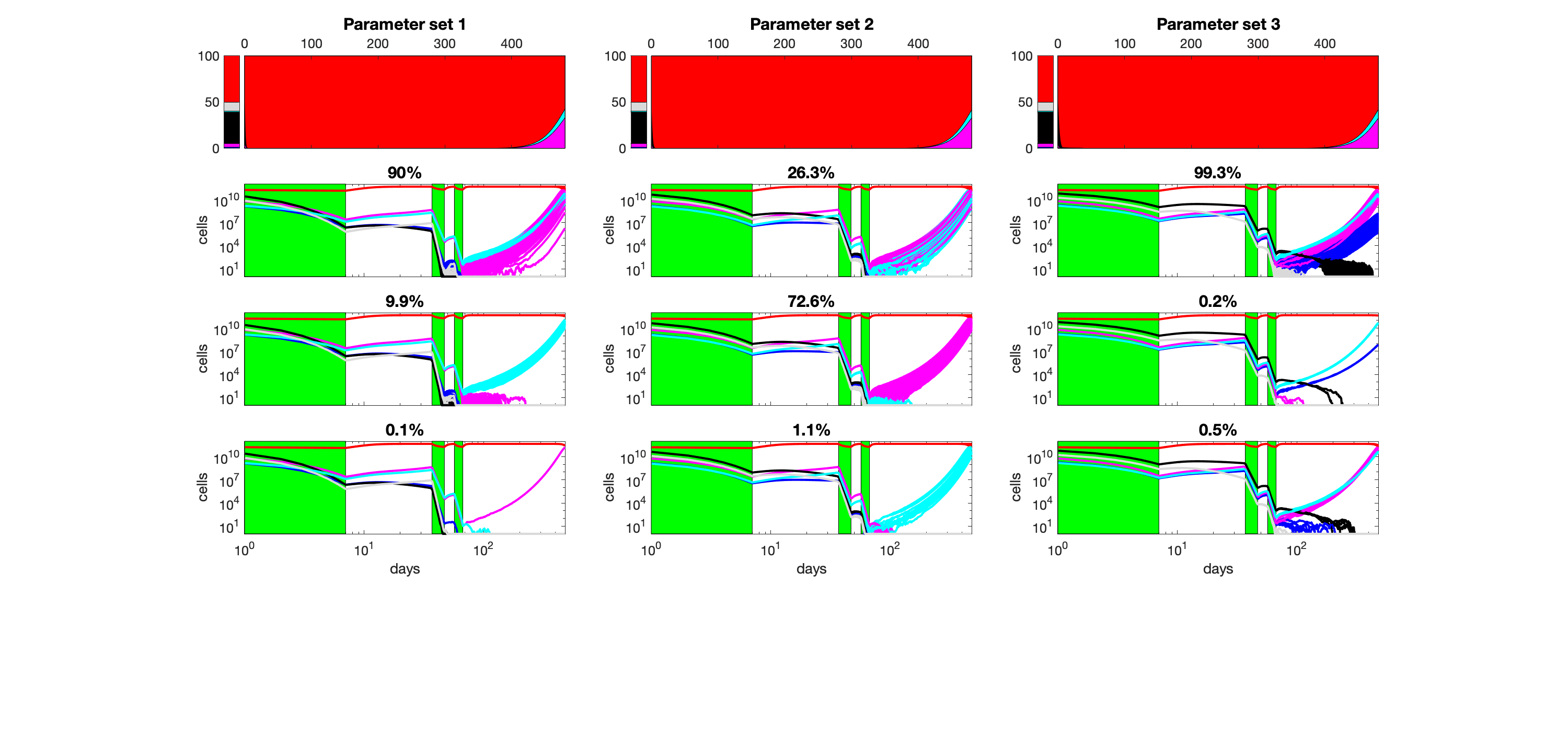

### S23_Fig.png

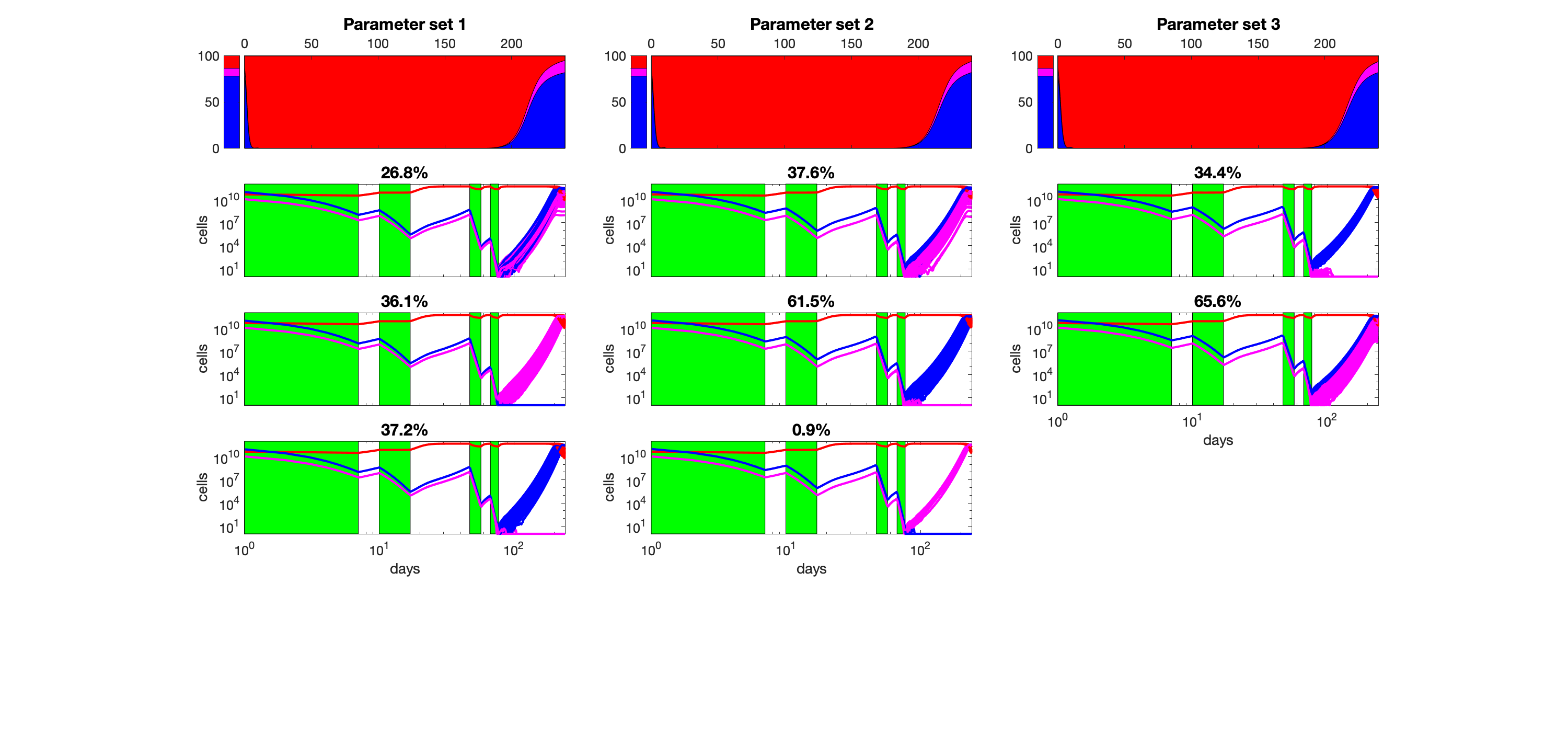

### S24_Fig.png

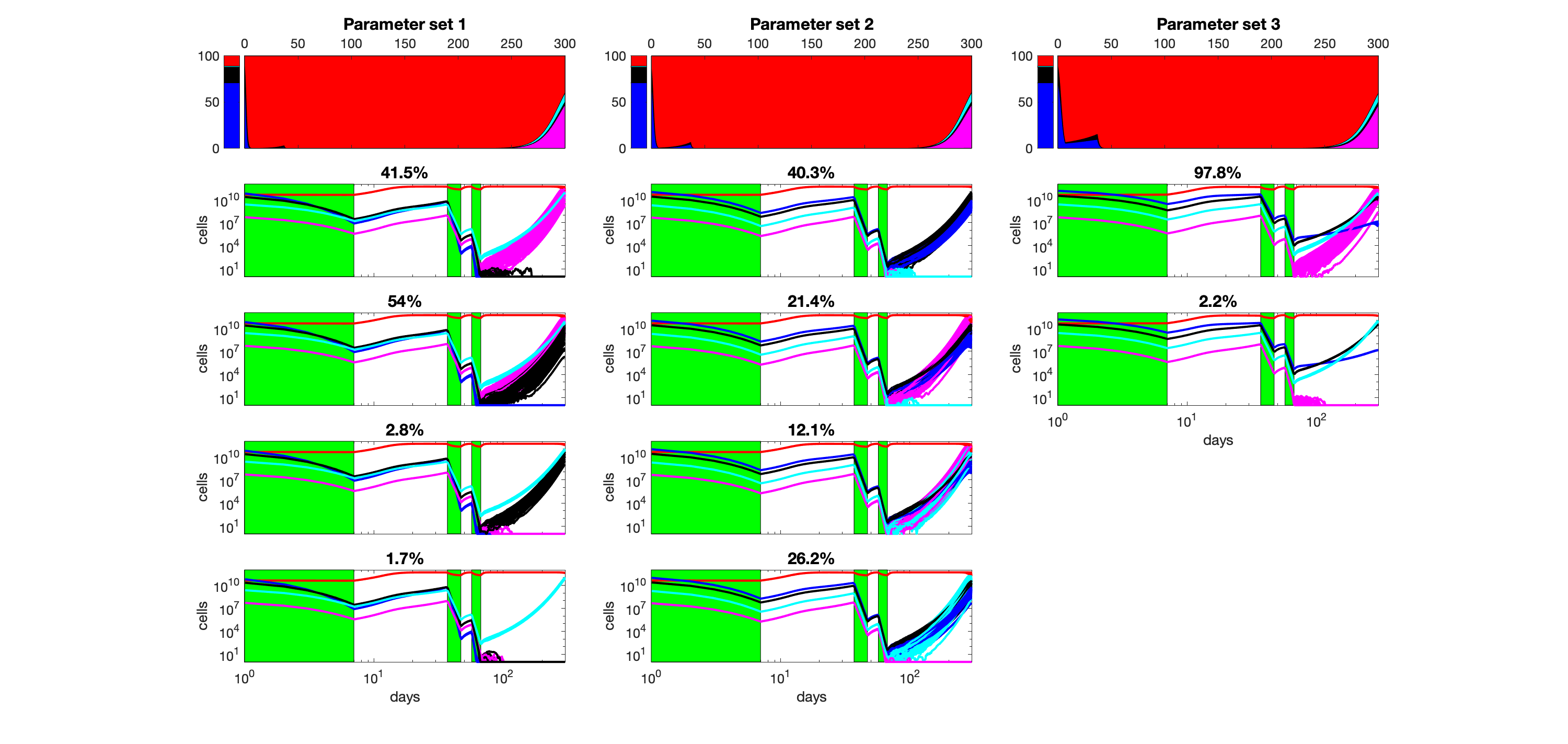
